# Knockout of Cyp26a1 and Cyp26b1 during post-natal life causes reduced lifespan, dermatitis, splenomegaly and systemic inflammation in mice

**DOI:** 10.1101/2020.07.09.196170

**Authors:** Jessica M Snyder, Guo Zhong, Cathryn Hogarth, Weize Huang, Traci Topping, Jeffrey LaFrance, Laura Palau, Lindsay C Czuba, Michael Griswold, Gabriel Ghiaur, Nina Isoherranen

## Abstract

All-*trans*-retinoic acid (*at*RA), the active metabolite of vitamin A, is an essential signaling molecule. Global knockout of the *at*RA clearing enzymes Cyp26a1 or Cyp26b1 is embryonic lethal. In adults, inhibition of Cyp26a1 and Cyp26b1 increases *at*RA concentrations and signaling. However, post-natal knockout of Cyp26a1 does not cause a severe phenotype. We hypothesized that Cyp26b1 is the main *at*RA clearing Cyp in post-natal mammals. This hypothesis was tested by generating tamoxifen inducible knockout mouse models of Cyp26b1 alone or with Cyp26a1. Both mouse models showed dermatitis, blepharitis and splenomegaly. Histology showed infiltration of inflammatory cells including neutrophils and T-lymphocytes into the skin and hyperkeratosis/hyperplasia of the non-glandular stomach. The mice lacking both Cyp26a1 and Cyp26b1 also failed to gain weight and showed fat atrophy. There were significant changes in vitamin A homeostasis demonstrating the paramount role of Cyp26b1 in regulating retinoid homeostasis in post-natal life.

## Introduction

Homeostasis and adequate dietary intake of the fat-soluble vitamin A (retinol) is critical for maintenance of health in humans and other mammals (Blomhoff and Blomhoff, 2006; Sommer, 2008; Gudas, 2012; O’Byrne and Blaner, 2013). In particular, the critical roles of the active metabolite of vitamin A, all-*trans*-retinoic acid (*at*RA), during fetal development, in reproduction, in the maintenance of healthy skin, epithelia, and immune system and in regulating cell division and hematopoiesis have been well characterized (Clagett-Dame and Knutson, 2011; Hall *et al.*, 2011; Rhinn and Dolle, 2012; Ghiaur *et al.*, 2013; Hogarth and Griswold, 2013; Raverdeau and Mills, 2014; Alonso *et al.*, 2017). Historically, analysis of the role of vitamin A and retinoid signaling has been focused on the effects of vitamin A deficiency on overall health (Sommer, 2008). However, distinct descriptions of hypervitaminosis A also exist, for example in arctic travelers (Biesalski, 1989; Senoo *et al.*, 2012). Hypervitaminosis A cases have changes in central nervous system, liver disorders, changes to the bone and effects on skin and mucous membranes, but the exact mechanisms and the retinoids causing these toxicities cannot be easily delineated (Biesalski, 1989). Chronic (13-week) dosing of *at*RA to rats (14 or 50 mg/kg) caused lymphoid hyperplasia, splenic extramedullary hematopoiesis, hyperkeratosis of the non-glandular stomach, long-bone fractures, hematological changes and testicular degeneration and atrophy (Kurtz *et al.*, 1984). After acute treatment the toxic doses are much higher (Kretzschmar and Leuschner, 1975; Biesalski, 1989). Treatment with 215 mg/kg po to mice or 400 mg/kg po to rats for five days led to death in 20% of the animals while lower doses caused characteristic symptoms of hair loss, dermal and mucosal alterations and loss of body weight (Kretzschmar and Leuschner, 1975; Biesalski, 1989). Genetic deficiencies in human *at*RA metabolism have supported the notion that excess endogenous *at*RA affects bone homeostasis and development (Laue *et al.*, 2011; Wen *et al.*, 2013; Nilsson *et al.*, 2016) and may affect skin (Slavotinek *et al.*, 2013) but limited information is available overall on the impact of genetic defects in *at*RA elimination. This is likely due to the critical function of *at*RA homeostasis in healthy embryogenesis and fetal development and incompatibility of genetic mutations with development (Isoherranen and Zhong, 2019).

The overall enzymology of vitamin A metabolism in mammals has been relatively well characterized (Napoli, 2012; Kedishvili, 2013; Li *et al.*, 2014; Isoherranen and Zhong, 2019). Vitamin A storage as retinyl esters in the liver stellate cells is mediated by LRAT, and the vitamin A stores are mobilized as retinol bound to the retinol binding protein RBP4. *at*RA is synthesized via a two-step process by retinol dehydrogenases and aldehyde dehydrogenases, and clearance of *at*RA is mediated by the Cyp26 enzymes. Knockout mouse models have been extensively used to define the role of the individual enzymes in regulating *at*RA gradients and signaling during development and organogenesis, but many of these knockout models are not viable (Duester, 2008; Rhinn and Dolle, 2012). Therefore, the role of specific enzymes in *at*RA homeostasis in different tissues during post-natal life is still not well understood.

The enzymes of the Cytochrome P450 family 26 (Cyp26) are known as the key *at*RA hydroxylases in all chordates (Isoherranen and Zhong, 2019). The Cyp26 family includes three enzymes, Cyp26a1, Cyp26b1, and Cyp26c1 of which Cyp26a1 and Cyp26b1 are necessary for embryonic development and survival during fetal life (White *et al.*, 1994, 2000; Fujii *et al.*, 1997; Taimi *et al.*, 2004; Pennimpede *et al.*, 2010). While Cyp26a1 knockout mice show a severe phenotype during fetal development (Rhinn and Dolle, 2012), global knockout of Cyp26a1 in juvenile or adult mice was recently shown not to cause classic retinoid toxicity or affect mouse health and survival significantly (Zhong *et al.*, 2019). This finding was surprising in light of the phenotype of Cyp26a1 embryonic knockout mice, and the data that show that inhibition of the Cyp26 enzymes increases tissue *at*RA concentrations and retinoid signaling (Stoppie *et al.*, 2000; Nelson *et al.*, 2013; Stevison *et al.*, 2017). In adult mice and humans, Cyp26a1 has been shown to be the predominant Cyp26 enzyme in the liver (Thatcher *et al.*, 2010), and Cyp26a1 appears to be important for clearing exogenous *at*RA (Zhong *et al.*, 2019). In contrast, our previous studies have suggested that Cyp26b1 is responsible for extrahepatic clearance of *at*RA in adult animals and humans, and that Cyp26b1 may be the critical enzyme regulating endogenous *at*RA signaling in adult animals (Topletz *et al.*, 2012; Stevison *et al.*, 2017; Zhong *et al.*, 2019; Hernandez *et al.*, 2020).

Based on the existing data, we hypothesized that post-natal inducible global knockout of Cyp26b1 or both Cyp26a1 and Cyp26b1 in adult or juvenile mice would result in decreased clearance of *at*RA, and therefore a significant increase in tissue *at*RA concentrations. We also predicted that such increased *at*RA concentrations would result in increased retinoid signaling in retinoid target organs and subsequently retinoid related toxicity. Given our previous data showing minimal phenotypic abnormalities after inducible global knockout of Cyp26a1 in juvenile and adult mice, and the modeling of critical role of Cyp26b1 in endogenous *at*RA clearance (Stevison *et al.*, 2017; Zhong *et al.*, 2019), we predicted that postnatal knockout of Cyp26b1 alone would lead to significantly decreased *at*RA clearance and retinoid toxicity. These hypotheses were also supported by the observed phenotype of testis cell type specific knockouts of Cyp26b1 and the T-cell specific knockout of Cyp26b1 (Takeuchi *et al.*, 2011; Hogarth *et al.*, 2015). These hypotheses were tested using a tamoxifen inducible Cyp26a1/Cyp26b1 Cre-Lox knockout mouse model with the aim to determine the importance of the Cyp26 enzymes in *at*RA clearance and in regulating *at*RA signaling in post-natal (juvenile and adult) mice.

## Methods

### Ethics statement, animal care and breeding

All the procedures involving mice prior to the commencement of these studies were approved by the Washington State University Committee on the Use and Care of Animals. The mouse colonies were maintained in a temperature- and humidity-controlled environment with food and water provided *ad libitum*. The mice were fed LabDiet 5K67 containing 1.5 ppm of carotene and 20 IU/g of vitamin A. The following mouse strains were used in these studies: The tamoxifen-inducible Cre recombinase line B6.129-*Gt(ROSA)26Sor*^*tm1(cre/ERT2)Tyjl*^*J* (stock number: 008463), herein referred to as Tam-Cre mice, purchased from JAX Mice (Bar Harbor, Maine, USA), and the *Cyp26a1*^*tm1Ptk*^ *(Cyp26a1)* and *Cyp26b1*^*tm1Ptk*^ *(Cyp26b1)* floxed mouse lines originally generated by Dr. Martin Petkovitch (Abu-Abed *et al.*, 2001) and reported previously (Hogarth *et al.*, 2015; Zhong *et al.*, 2019). *Cyp26a1* and *Cyp26b1* floxed mice were bred to Tam-Cre-positive mice using a standard Cre-Lox breeding scheme and the following four genotypes were analyzed: 1) homozygous flox *Cyp26a1/b1*/Tam-Cre-negative (controls, *Cyp26a1*^*fl/fl*^*b1*^*fl/fl*^, referred to as Cyp26a1^+/+^b1^+/+^), 2) homozygous floxed *Cyp26a1* and *Cyp26b1*/Tam-Cre-positive (double knock outs, *Cyp26a1*^*fl/fl*^*b1*^*fl/fl*^, referred to as Cyp26a1^−/−^b1^−/−^). 3) *Cyp26a1* wild type and heterozygous flox *Cyp26b1*/Tam-Cre-positive (*Cyp26b1* heterozygous knock outs, *Cyp26b1*^*wt/fl*^, referred to as Cyp26b1^+/−^), and 4) *Cyp26a1* wild type and homozygous flox *Cyp26b1*/Tam-Cre-positive (*Cyp26b1* knock outs, *Cyp26b1*^*fl/fl*^, referred to as Cyp26b1^−/−^). All four groups were treated with tamoxifen to induce expression of Cre recombinase, if present, as described below. The day of birth was designated as day 0.

The effect of the genotypes was studied in four separate cohorts of mice. First cohort (cohort 1) included control and Cyp26a1^−/−^b1^−/−^ mice with tamoxifen treatments as juveniles or adults, second cohort (cohort 2) included Cyp26b1^+/−^ and Cyp26b1^−/−^ mice with tamoxifen treatments as juveniles or adults and the third cohort (cohort 3) was a repeated breeding of control and Cyp26a1^−/−^b1^−/−^ mice with tamoxifen treatment as juvenile mice. An additional cohort of control and Cyp26a1^−/−^b1^−/−^ (cohort 4), mice was bred at the Johns Hopkins university for the pharmacokinetic (PK) studies and these studies were approved by the animal care and use committee at Johns Hopkins University Animal Facility. For these studies female Cyp26a1^−/−^b1^−/−^ and control mice were weaned at 3 weeks of age, and recombination was induced at 4 weeks of age with tamoxifen injections as described above and as previously published (Zhong *et al.*, 2019). These mice were fed a Teklad Global 18% Protein-extruded rodent diet (Teklad Diets, Madison WI) containing 30 IU/g of vitamin A.

Animals were genotyped and knockouts assessed by PCR using the following primer sets: for Tam-Cre forward primer GCATTACCGGTCGATGCAACGAGTG and reverse primer GAACGCTAGAGCCTGTTTTGCACGTTC; for Cyp26a1 Flox forward primer ACATTGCAGATGGTGCTTCA and reverse primer CGTATTTCCTGCGCTTCATC; for Cyp26b1 Flox forward primer CAGTAGATGTTTGAGTGACACAGCC and reverse primer GAGGAAGTGTCAGGAGAAGTGG. In addition mRNA was extracted from livers of the experimental animals and controls and the expression of Cyp26a1 mRNA and other relevant genes was analyzed by q-RT-PCR as previously described (Zhong *et al.*, 2019).

Tamoxifen (Sigma-Aldrich, St Louis, MI T5648) was dissolved in 10% ethanol and 90% sesame oil at a concentration of 20 mg/ml, and the solution was wrapped in foil to protect it from light. Mice were intraperitoneally (ip) injected with 80 mg/kg tamoxifen once a day for 5 consecutive days beginning at 21 days post-partum (dpp) for the juvenile induction studies or 60 dpp for the adult induction studies. The animals were then left to recover for 8 weeks after the final tamoxifen injection before being euthanized for tissue collection, unless they met euthanasia criteria (detailed below) earlier than 8 weeks. Animal health was monitored via assessment of weight twice a week after tamoxifen injection. Animals were euthanized by CO_2_ asphyxiation followed by cervical dislocation. Blood was collected by cardiac puncture immediately following CO_2_ asphyxiation and cervical dislocation. Major organs were collected and organ weights were measured at necropsy for adult (Cyp26a1^−/−^b1^−/−^ n= 11, 5 males [M] and 6 females [F]; Cyp26b1^−/−^ n= 9, 5M, 4F; Cyp26b1^+/−^ n= 10, 5M, 5F; control n = 10, 5M, 5F) and juvenile (Cyp26a1^−/−^b1^−/−^ n= 29, 15M, 14F; Cyp26b1^−/−^ n= 15, 6M, 9F; Cyp26b1^+/−^ n= 10, 5M, 5F; control n= 22, 12M, 10F) induction mice.

### Body weight and survival analysis

Based on the poor health of the very first Cyp26b1^−/−^ and Cyp26a1^−/−^b1^−/−^ animals generated following tamoxifen injection, a monitoring protocol and euthanasia criteria were established with assistance from the Washington State University Office of the Campus Veterinarian. The monitoring protocol was as follows: at the first sign of ill health, most often being dermatitis lesions on the ears, the animals were flagged as needing a daily health check involving weight measurement, treatment, and body condition scoring (Ullman-Culleré and Foltz, 1999). An animal was euthanized if they had lost more than 15% of their pre-tamoxifen injection body weight or received a body condition score of 2 or less regardless of the timing following tamoxifen injection. Animal necropsy and tissue collection was performed as described below. During the monitoring process, any dermatitis lesions were treated with an antibiotic ointment free of retinoids (Equate Antibiotic + Pain Relief NCD 49035-840-01) to prevent infection and mice were given a daily 1cm^3^ piece of Nutrigel nutritional supplement (Nutra-Gel Diet™, # S4798, Bio-Serve, Flemington, NJ with Vitamin A at 5519 IU/kg).

### Histological analyses

For all histological analyses, mice were euthanized and subjected to routine necropsy (Treuting and Snyder, 2015). Mouse tissues were immersed in 10% neutral-buffered formalin for 48 hours and then stored in 70% ethanol until routine processing in which they were embedded in paraffin, cut into 4-5 μm sections and hematoxylin and eosin stained. To study the impact of induction of *Cyp26a1* and *Cyp26b1* knock out in adulthood, 6 controls (3M, 3F); 6 Cyp26a1^−/−^b1^−/−^ (3M, 3F); 7 Cyp26b1^−/−^ (3M, 4F), and 6 Cyp26b1^+/−^ (3M, 3F) mice were examined histologically following tamoxifen administration at 60 dpp (adults). To characterize the impact of *Cyp26a1* and *Cyp26b1* knock-out in juvenile animals, 6 controls (3M, 3F), 7 Cyp26a1^−/−^b1^−/−^ (3M, 4F); 7 Cyp26b1^−/−^ (3M, 4F), and 6 Cyp26b1^+/−^ (3M, 3F) mice were examined histologically following tamoxifen administration at 21 dpp. For both groups, necropsies were performed 60 days following the last tamoxifen injection except one Cyp26a1^−/−^b1^−/−^ from the adult induction group that met euthanasia criteria and was sacrificed early, at 6 weeks, and two Cyp26a1^−/−^b1^−/−^ from the juvenile induction group that were sacrificed at 21 days following the last tamoxifen injection. Tissues examined on initial phenotyping included: decalcified cross section of the skull including brain and bulla; lung; thymus; heart; kidney; liver; pancreas and mesentery; lymph node; spleen; reproductive tract; bladder; gastrointestinal tract including esophagus, stomach, small intestine and large intestine; ear pinna; and intrascapular skin. Following initial review of the tissues, a targeted organ histology including decalcified cross section skull; thymus; pancreas and mesentery; spleen; lymph node; reproductive tract; decalcified cross section of the stifle with distal femur, proximal tibia, and associated skeletal muscle; esophagus; stomach; skin and pinna, was performed in subsequent animals.

Lesions including dermatitis of the skin of the dorsal head and muzzle, interscapular region, and ear pinna; blepharitis; keratin accumulation in the external ear canal; gastritis; hyperkeratosis and hyperplasia of the non-glandular stomach; testicular degeneration; expansion of the splenic red pulp by hematopoietic cells including myeloid and erythroid precursors and megakaryocytes (interpreted as increased extramedullary hematopoiesis); and lymph node hyperplasia were scored on a scale of 0-4, with 0 representing normal, 1 representing minimal, 2 representing mild, 3 representing moderate, and 4 representing severe histological changes. Additional details on the scoring system are included in supplemental material. Tissues were evaluated and scored by a board-certified veterinary pathologist who was not blinded to genotype and experimental manipulation for the initial group of juvenile animals evaluated during development of the scoring system, but who was blinded to genotype for remaining juvenile animals (3 control mice, 4 Cyp26b1^+/−^ mice, and 4 Cyp26b1^−/−^ mice) and for all adult animals. Juvenile mice evaluated initially in an unblinded fashion were re-scored blindly prior to manuscript preparation. Representative images were taken using NIS-Elements BR 3.2 64-bit and plated in Adobe Photoshop Elements. Image white balance, brightness and contrast were adjusted using auto corrections applied to the entire image. Original magnification is stated.

Immunohistochemistry, tissue processing and staining were done by the University of Washington Histology and Imaging core, Department of Comparative Medicine on formalin-fixed paraffin embedded sections using the following primary antibodies: rat monoclonal anti F4/80 (Clone BM8, Invitrogen, Cat. No. MF48000, 1:200 in Bond Primary Antibody Diluent); rat monoclonal anti CD45 (Clone RA3-6B2, BD Pharmingen, Cat. No. 557390,1:3000 dilution); rabbit monoclonal anti Ki67 (Clone D3B5, Cell Signaling, Cat. No. 12202, 1:400 dilution); rat monoclonal anti CD3 (Clone CD3-12, Abd Serotec, Cat. No. MCA1477, 1:100 dilution); goat polyclonal anti YM1/Chitinase-3 like 3 (R&D systems, Cat. No. AF2446, 1:400 dilution). Antigen retrieval of HIER2 (EDTA) for 10 (Ki67) or 20 (CD3) minutes at 100°C; HIER1 (Citrate) for 10 minutes at 100°C (CD45); or PK15 at 37°C (F4/80); or no antigen retrieval (YM1) was used. Slides were run on a Bond autostainer platform. The primary antibodies were detected using the Bond Polymer Refine (DAB) Detection Kit (Leica Biosystems; Buffalo Grove, IL). Positive control mouse tissues for all antibodies were run in addition to negative controls performed using rat or rabbit isotype IgG.

Immunohistochemistry for CD3+ (T lymphocytes); CD45+ (B lymphocytes); and F4/80 (macrophages) was performed on pinna and immunohistochemistry for Ki67 was performed on the head from one male and one female of Cyp26a1^+/+^b1^+/+^ and one male and one female of Cyp26a1^−/−^b1^−/−^ juvenile-induction mice. Immunohistochemistry for Ki67 and CD3 was also performed on the stomach and small intestine from 3 female Cyp26a1^−/−^b1^−/−^, 3 female Cyp26a1^+/+^b1^−/−^, and 2 female Cyp26a1^+/+^b1^+/+^ juvenile-induction animals. Immunohistochemistry for YM1 was performed on decalcified cross section of skull (nasal cavity) and stomach from one male Cyp26a1^−/−^b1^−/−^ mouse. Special stains including Toluidine Blue (to highlight mast cells), and Masson’s trichrome (to highlight collagen) were also performed on skin from one male and one female Cyp26a1^+/+^b1^+/+^ and one male and one female Cyp26a1^−/−^b1^−/−^ juvenile-induction mice.

### Quantitative IHC

Image analysis on the small intestine (CD3) and stomach (Ki67) was performed using whole slide digital images and automated image analysis. All slides were scanned in bright field with a 20X objective using a Nanozoomer Digital Pathology slide scanner (Hamamatsu; Bridgewater, New Jersey). Whole slide digital images were imported into Visiopharm software (Hoersholm, Denmark) for analysis. The software converted the initial digital imaging into gray scale values using two features, RGB-R with a mean filter of 5 pixels by 5 pixels and an RGB-B feature. Visiopharm was then trained to label positive staining and the background tissue counter stain using a project-specific configuration based on threshold pixel values. Images were processed in batch mode using this configuration to generate the desired outputs (ex. area of CD3 and ratio of CD3 to total tissue area).

### Quantification of atRA, retinol and retinyl esters in mouse serum and tissues by LC-MS/MS

The concentrations of retinyl palmitate (RE), retinol (ROL) and *at*RA in mouse serum, liver, testis and spleen were measured as previously described (Zhong *et al.*, 2019). Retinoid concentrations in small intestine (130-160 mg) and skin (100-130 mg) were measured using the same protocol as used for the measurement of spleen retinoids (Zhong *et al.*, 2019). Spleen, skin and small intestine *at*RA samples were analyzed using AB Sciex 5500 QTRAP mass spectrometer (AB Sciex LLC; Framingham, MA) coupled with an Agilent 1290 UHPLC (Agilent Technologies; Santa Clara, CA) while all other retinoid samples were analyzed using AB Sciex 6500 QTRAP mass spectrometer coupled with a Shimadzu UFLC XR DGU-20A5 (Shimadzu Corp.; Kyoto, Japan). The LC-MS/MS methods were the same as described previously (Zhong *et al.*, 2019) and all the data were analyzed using Analyst software by two independent individuals of whom one was blinded to the genotype of the animals in the analyzed samples.

### Quantification of RBP4 and IL-17 in serum and liver using ELISA

The RBP4 concentrations in mouse serum and liver were quantified using a mouse RBP4 ELISA kit (ab202404; Abcam; Cambridge, UK) according to the manufacturer’s instructions. Mouse serum samples were diluted with the sample diluent buffer provided in the kit in a ratio of 1 to 200,000. Mouse liver tissues of about 50 mg were homogenized with 5x tissue weight of cell extraction buffer provided by the kit. The liver homogenates were centrifuged at 18,000 g at 4°C for 20 minutes and the supernatant was collected. The protein concentrations of the supernatant were determined using the bicinchoninic acid (BCA) assay (Thermo Fisher Scientific, Waltham, MA) and were diluted to 2.5 μg/mL with the cell extraction buffer. Diluted serum and liver samples were used to measure the serum and liver RBP4 concentrations following the procedures as described previously (Zhong *et al.*, 2019).

The serum IL-17 concentrations were quantified using a mouse IL-17 ELISA kit from Abcam (ab100702) according to the manufacturer’s instructions. Briefly, mouse serum samples were diluted with the dilution buffer provided by the kit in a ratio of 1 to 3. Diluted serum and calibration standards were first incubated in the ELISA microplate for 2.5 hr with shaking at 4,000 rpm at room temperature. After the plate was washed four times with washing buffer, the IL-17 detection antibody was added and the ELISA plate was incubated at room temperature for 1 hr with gentle shaking. After the plate was washed for four times, the horseradish peroxidase (HRP)-streptavidin solution was added and the plate was incubated at room temperature for 45 minutes. After the plate was washed, 3,3,5,5-tetramethylbenzidine substrate was added and incubated for 30 min followed by the addition of the stop solution. Absorbance was measured at 450 nm immediately, and IL-17 concentrations were calculated using the calibration curve provided by the kit.

### Pharmacokinetics of exogenously administered atRA

To study the impact of loss of Cyp26a1 and Cyp26b1 activity to exogenous *at*RA PK, mice were administered one dose of 1 mg/kg *at*RA ip at six weeks of age (2 weeks post-tamoxifen injection). Blood was collected at 0 h, 0.5 h, 1 h, 2 h, 4 h and 6 h and plasma was separated by centrifugation. Throughout the plasma collection period the mice were observed for signs of acute toxicity. *at*RA concentrations were analyzed by LC-MS/MS as previously described (Zhong *et al.*, 2019). The pharmacokinetics of *at*RA were analyzed using Phoenix software (Certara, Princeton, NJ) and standard noncompartmental analysis. Maximum concentration (*C*_max_), area under the plasma concentration-time curve from time 0 to infinity (AUC0_→∞_), clearance (CL), and half-life (*t*_½_) values were calculated. A separate group of Cyp26a1^−/−^b1^−/−^ and Cyp26a1/b1^+/+^ mice at 7 weeks post tamoxifen injection was injected with 10 mg/kg *at*RA ip as described previously (Zhong *et al.*, 2019). These animals were intended for a PK study but at 30 minutes after *at*RA administration, Cyp26a1^−/−^b1^−/−^ mice started to present lethargy. At 2h after *at*RA administration, 90% of Cyp26 deficient mice had died while all Cyp26a1^+/+^b1^+/+^ mice were alive.

### Statistical analyses

The impact of Cyp26 genotype and sex on the mouse body weight gain per day and the mouse body weight at baseline (t=0) was evaluated by a multivariate linear mixed effect model using R (R Foundation for Statistical Computing, Vienna, Austria) as previously described (Zhong *et al.*, 2019). In brief, the multivariate linear mixed effect model was as follows: Body weight (kg) = a1 + b1×time + b2×gene1 + b3×gene2 + b4×sex + c1×time×gene1 + c2×time×gene2 + c3×sex×gene1 + c4×sex×gene2 + c5×sex×time + c6×sex×time×gene1 + c7×sex×time×gene2 where time is treated as continuous variable with each value indicating the number of days after the tamoxifen treatment; gene1 and gene2 are binary variables for genotype (either Cyp26b1^+/+^, Cyp26b1^−/+^, Cyp26b1^−/−^, or Cyp26a1^+/+^b1^+/+^, Cyp26a1^+/−^b1^−/+^, Cyp26a1^−/−^b1^−/−^). Sex is a binary variable for female (0) and male (1). This modeling framework enables robust statistical inference of Cyp26 genotype and sex effects on the mouse body weight gain (slopes) and the mouse body weight at baseline (intercepts). Each coefficient was tested to evaluate whether its value is statistically significantly different from zero. The inference test of coefficient linear combination was conducted when significance of the sum of two coefficients was queried. A nominal value of 0.05 was considered significant with Bonferroni correction made for multiple comparisons.

The differences of retinoid (retinyl palmitate, retinol, *at*RA) and RBP4 concentrations in serum and tissues (spleen, liver, skin) and in terminal body weight normalize spleen weights among the experimental groups with different sex and Cyp26 genotypes was evaluated by a multivariate linear regression using R. The multivariate linear regression treats retinoids and RBP4 concentrations as response variables, Cyp26 genotype and sex as two explanatory variables, and incorporates the interaction effect between Cyp26 genotype and sex. This modeling framework allows robust statistical inference of Cyp26 genotype and sex effects on the retinoids and RBP4 concentrations in serum and multiple tissues in mouse. The multivariate linear regression model was as follows: Retinoids and RBP4 concentrations in serum and tissues = a1 + b1×sex + b2×gene + b3×sex×gene where sex is a binary variable for female (0) and male (1) and gene is a binary variable for genotype (either Cyp26b1^+/−^ vs Cyp26b1^−/−^, or Cyp26a1^+/+^b1^+/+^ vs Cyp26a1^−/−^b1^−/−^). Each coefficient was tested to evaluate whether its value is statistically significantly different from zero. A nominal value of 0.05 was considered significant with Bonferroni correction made for multiple comparisons. For replicate mouse cohort studies (Cyp26a1^−/−^b1^−/−^ juvenile induction studies) findings were considered significant only if they were reproduced in both cohorts of mice. The differences between Cyp26a1^−/−^b1^−/−^ mice versus controls for semiquantitative histological analyses and serum concentrations of IL-17 was assessed by Mann-Whitney test.

## Results

### Post-natal global deletion of Cyp26a1 and Cyp26b1 severely compromises animal health

Following a standard Cre-Lox breeding scheme, mice homozygous for floxed alleles of both *Cyp26a1* and *Cyp26b1* or *Cyp26b1* alone and carrying the tamoxifen cre transgene were successfully generated. Correct LoxP excision to generate the mutant alleles following tamoxifen injections was confirmed via PCR analysis. The knock-out was induced in either juvenile (21 days old) or adult mice (60 days old) to explore possible differences in the role of these enzymes in retinoid homeostasis and signaling as mice age. Based on mRNA analysis of the livers, *Cyp26a1* was efficiently deleted in the Cyp26a1^−/−^b1^−/−^ mice (Supplemental Figure 1). Gross phenotypic monitoring of Cyp26a1^−/−^b1^−/−^ and Cyp26b1^−/−^ mice revealed severe dermatitis and blepharitis which began approximately 10-14 days after the last tamoxifen injection. For humane reasons, treatment with antibiotic ointment was initiated when mild to moderate dermatitis was identified to prevent infection, and the animals were offered a nutritional supplement that could be easily chewed and swallowed on a daily basis. This did not stop the progression of clinical illness. Of the Cyp26a1^−/−^b1^−/−^ mice in which the knockout was induced as juveniles, only 54.5% of males and 59.1% of females survived till the end of the 60-day observation period (Figure 1). When the Cyp26a1^−/−^b1^−/−^ knockout was induced in adults, 55.6% of female and 100% of male mice survived through the observation period (Figure 1). The body weight gain was significantly lower in all groups of Cyp26a1^−/−^b1^−/−^ mice (Figure 1, Supplemental Figure 2). In contrast to the Cyp26a1^−/−^b1^−/−^ mice, Cyp26b1^−/−^ mice all survived through the 60-day observation period despite developing dermatitis. The body weight in male adult-induction Cyp26b1^−/−^ mice was significantly lower than controls while in females and in male Cyp26b1^−/−^ juvenile-induction mice, there was no difference in body weights compared to controls. Abnormalities on gross examination of the Cyp26a1^−/−^b1^−/−^ and the Cyp26b1^−/−^ mice included a moderate to severe blepharitis, dermatitis that was most pronounced on the dorsum of the head and the ear pinna and a gray color to the fur around the eyes and face. These abnormalities were not seen in any of the control mice examined.

**Figure 1.**
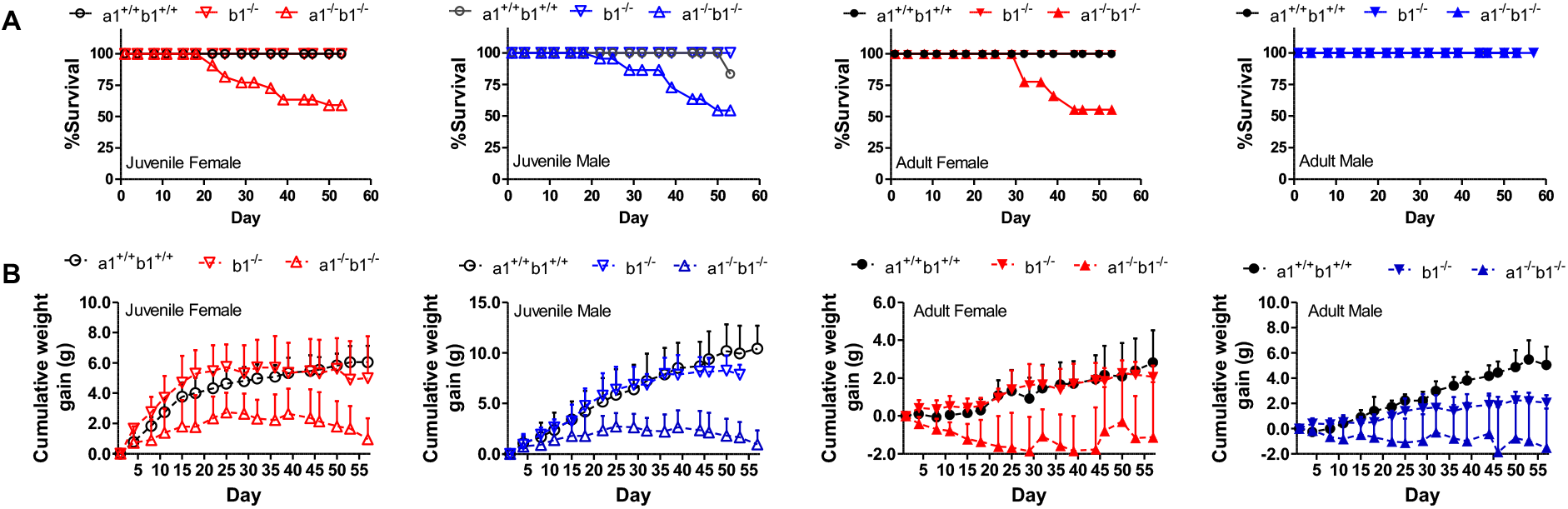
Survival curves (A) and cumulative weight gain (B) in control (a1^+/+^b1^+/+^, Cyp26a1^+/+^b1^+/+^), Cyp26b1^−/−^ (b1^−/−^) and Cyp26a1^−/−^b1^−/−^ (a1^−/−^b1^−/−^) mice. The x axis shows days after tamoxifen injection. Open and filled symbols represent juvenile and adult mice, respectively. Data points show mean values with standard deviation (SD) included as error bars. All the data are from mice in cohorts 1 and 2. The data for all individual mice and for all experimental groups are shown in Supplemental Figure 2.

### Vitamin A metabolome is altered in the serum and liver of Cyp26a1^−/−^b1^−/−^ and Cyp26b1^−/−^ mice without histological differences in major organs

To explore the relationship between the severe phenotype of the Cyp26a1^−/−^b1^−/−^ and Cyp26b1^−/−^ mice and altered retinoid homeostasis, the effect of deletion of Cyp26a1 and Cyp26b1 activity on serum and liver vitamin A metabolome was studied. Histology of major organs was also conducted to characterize the phenotype. The serum retinyl esters, retinol, *at*RA and RBP4 concentrations were analyzed in all groups of mice studied. Significant changes in serum vitamin A metabolome were observed but the effects were dependent on genotype and the age of the mice when knockout was induced. The serum retinol and RBP4 concentrations were consistently decreased in the Cyp26a1^−/−^b1^−/−^ mice in which the knockout was induced as juveniles in comparison to controls (Figure 2 and Supplemental Figure 3), but there was no change in serum retinol and RBP4 in Cyp26a1^−/−^b1^−/−^ mice in which the knockout was induced as adults (Supplemental Figure 3). In contrast, serum retinol concentrations were significantly decreased in Cyp26b1^−/−^ mice when compared to Cyp26b1^+/−^ mice for both juvenile and adult induction mice (Figure 2 and Supplemental Figure 3). Despite the changes in serum retinol, RBP4 concentrations were unaltered in the Cyp26b1^−/−^ mice (Figure 2 and Supplemental Figure 3). Serum retinyl ester and *at*RA concentrations were similar across all mouse genotype groups within an age group of mice, but older mice appeared to have higher *at*RA serum concentrations than younger mice, a finding consistent with our previous studies in similar age mice (Zhong *et al.*, 2019).

**Figure 2.**
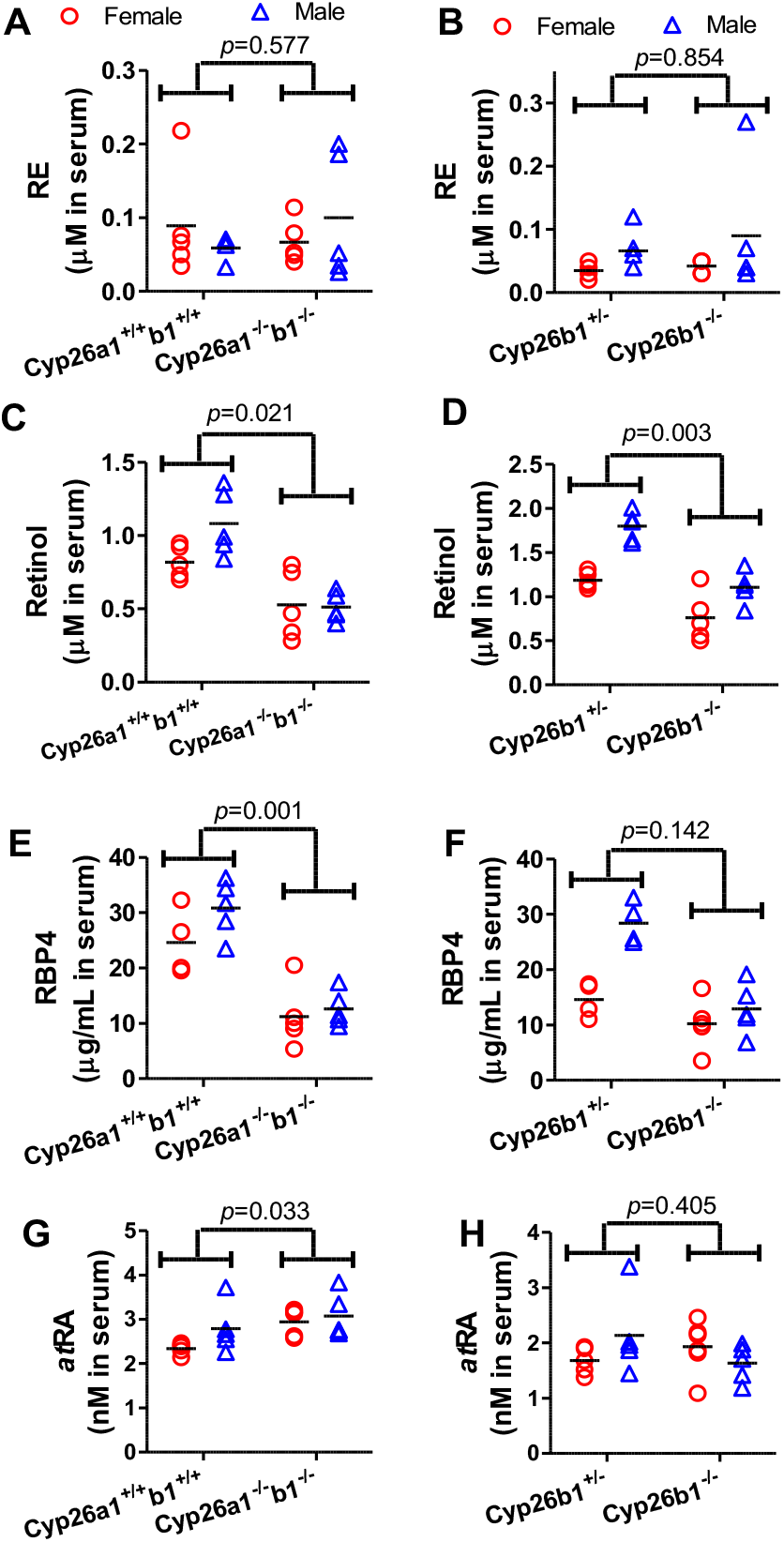
Serum retinyl palmitate (A and B), retinol (C and D), RBP4 (E and F) and atRA (G and H) concentrations in Cyp26a1^+/+^b1^+/+^ mice in comparison to Cyp26a1^−/−^b1^−/−^ mice (A, C, E and G) and in Cyp26b1^+/−^ mice in comparison to Cyp26b1^−/−^ mice (B, D, F and H). Each data point represents an individual mouse. The horizontal lines indicate mean values within the group. All the data shown is from juvenile mice from cohorts 2 and 3. Differences between groups were tested using multivariate linear regression as described in methods section and individual values are shown. The data for the adult mice and repeated juvenile mice are shown in Supplemental Figure 3.

No consistent histologic abnormalities of major organs including the brain, heart, lungs, pancreas, and kidney were observed in Cyp26a1^−/−^b1^−/−^ and Cyp26b1^−/−^ mice (data not shown). The livers of the Cyp26a1^−/−^b1^−/−^ and Cyp26b1^−/−^ mice generally appeared histologically normal although mild extramedullary proliferation of hematopoietic cells in Cyp26a1^−/−^b1^−/−^ (and minimal in occasional Cyp26b1^−/−^) mice was observed in the liver (Figure 3 and Supplemental Figure 7). Minimal multifocal hepatic necrosis and microgranulomas were also observed in some mice from all genotypes and considered incidental background lesions. The poor health status matching the lack of weight gain and weight loss in the Cyp26a1^−/−^b1^−/−^ and some of the Cyp26b1^−/−^ mice was also clearly observed in the histology of the abdominal white adipose tissue (Figure 3 and Supplemental Figure 4). Fat atrophy and smaller adipocytes were observed in the Cyp26a1^−/−^b1^−/−^ mice and to a lesser degree in the Cyp26b1^−/−^ mice when compared to controls (Figure 3 and Supplemental Figure 4). All mice, including controls, had minimal to moderate chronic to chronic-active inflammation of the mesenteric fat with low numbers of histiocytes with vacuolated cytoplasm, most likely related to intraperitoneal tamoxifen injection. However, only the juvenile induction Cyp26a1^−/−^b1^−/−^ mice, and not Cyp26b1^−/−^ mice or adult induction Cyp26a1^−/−^b1^−/−^ mice, had significantly higher mesenteric inflammation scores compared to controls, with a more pronounced neutrophilic component to the inflammation (Figure 3, Supplemental Figure 4).

**Figure 3.**
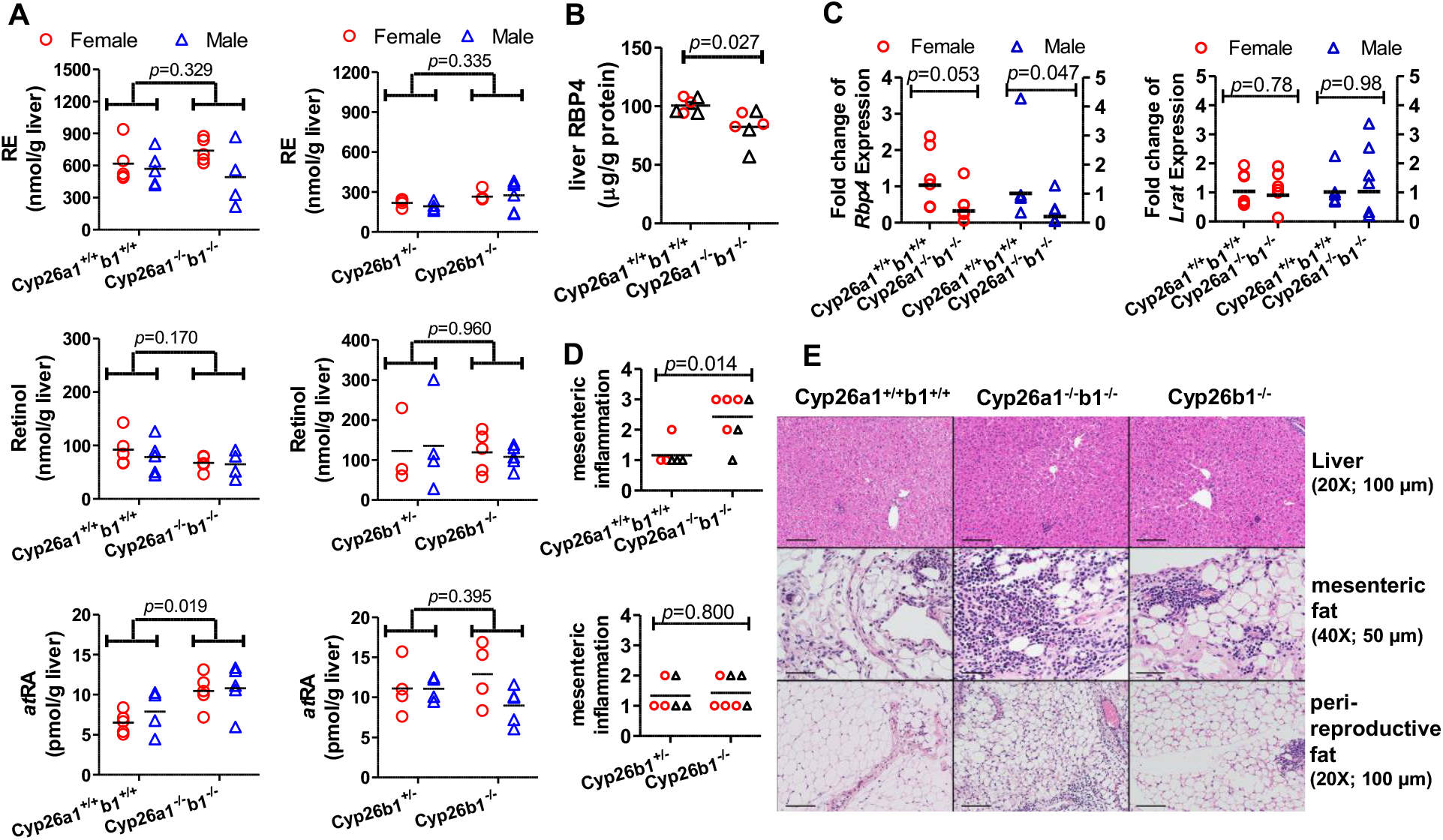
Liver retinyl palmitate, retinol and *at*RA concentrations (A), RBP4 expression level (B), liver *Rbp4* and *Lrat* mRNA expression (C), histological scores (D) and H&E stained representative histology images of liver and adipose tissues (E). The histology images are from the juvenile female induction mice in cohorts 1 and 2 (E). All the data in panels A-D are from the juvenile induction mice in cohorts 2 and 3. The horizontal lines indicate arithmetic mean values (A and B) and geometric mean values (C and D). In panel A, differences were tested by multivariate linear regression and obtained *p-*values are shown. To test for differences for RBP4 protein (B) the data were log transformed and analyzed by unpaired two-tailed *t*-test. In panel C, the fold change of mRNA expression was the average value calculated from two rt-PCR analyses performed on separate days. Female and male data were analyzed separately, and the data correspond to the left and right y axis, respectively. Data were log transformed and statistical significance was tested by unpaired two-tailed t-test. In panel D, statistical significance was tested by Mann-Whitney test. Each data point represents an individual mouse (A-D). In panels B and D open circles represent female mice and open triangles show male mice. In panel E, the original image magnification and the length the black scale bars are shown inside parentheses. The corresponding data for adult mice and a second cohort of juvenile induction mice are shown in Supplemental Figure 4.

In the liver, *at*RA concentrations were significantly increased in the Cyp26a1^−/−^b1^−/−^ mice in which the knock-out was induced as juveniles in comparison to control mice (Figure 3). This increase was reproduced in another group of juvenile induction mice (Supplemental Figure 4). However, no genotype effect on liver *at*RA concentrations was observed in the mice in which Cyp26a1^−/−^b1^−/−^ knock-out was induced as adults nor in any of the groups of Cyp26b1^−/−^ mice. No consistent changes in retinyl palmitate (RE) or retinol concentrations in the liver were detected across genotype groups (Figure 3, Supplemental Figure 4) consistent with the lack of change in *Lrat* mRNA in the liver (Figure 3). While retinol concentrations in the liver were largely unchanged, RBP4 protein expression was significantly decreased in the livers of Cyp26a1^−/−^b1^−/−^ mice when compared to control mice although the magnitude of change was small (Figure 3). This decrease in hepatic RBP4 expression appears to correlate with the decrease in serum retinol and possibly serum RBP4 rather than liver retinol concentrations, suggesting that synthesis of RBP4 in the liver may be regulating circulating retinol concentrations.

Consistent with our previous findings (Zhong *et al.*, 2019), sex differences were also detected in the retinoid concentrations in serum (Supplemental Table 1, Figure 2, Supplemental Figure 3). In all groups of mice, retinol concentrations in serum were significantly higher in male mice when compared to female mice (*p*-values listed in Supplemental Tables 1 and 2), and RBP4 concentrations were significantly higher or approached significance in male mice when compared to female mice (Supplemental Tables 1 and 2). In addition, serum retinyl palmitate concentrations were higher in adult male mice when compared to female mice, but no difference between male and female mice was observed in the juvenile induction mice (Supplemental Tables 1 and 2, Figure 2, Supplemental Figure 3). No sex differences were detected in the liver for retinyl palmitate or retinol (Figure 3, Supplemental Tables 1 and 2) but *at*RA concentrations were significantly higher in adult male mice than in adult female mice. No sex difference in *at*RA concentrations was detected in the liver of the younger mice or in the serum of any of the mouse groups studied (Supplemental Tables 1 and 2).

### Global deletion of Cyp26a1 and Cyp26b1 or Cyp26b1 alone is associated with severe dermatitis and blepharitis, with increased skin atRA concentrations and infiltration of inflammatory cells

To further characterize the observed skin lesions in the Cyp26a1^−/−^b1^−/−^ and Cyp26b1^−/−^ mice, skin histology was performed. Both Cyp26a1^−/−^b1^−/−^ and Cyp26b1^−/−^ mice showed moderate to marked dermatitis with epithelial hyperplasia/acanthosis, hyperkeratosis, and variable erosion and ulceration that was most pronounced on the dorsum of the head, ear pinnae and eyelids (blepharitis) (Figure 4, Figure 5 and Supplemental Figure 5). The inflammation varied in severity regionally and between mice and ranged from chronic to chronic-active to suppurative. Dermatitis and blepharitis were observed both in the mice in which the knockout was induced as juveniles and as adults, and significant increases in the scores for head dermatitis and blepharitis were observed in all groups of Cyp26a1^−/−^b1^−/−^ and Cyp26b1^−/−^ mice (Figure 4 and Supplemental Figure 5). The interscapular skin was less severely and more sporadically affected (Figure 4 and Supplemental Figure 5). All Cyp26a1^−/−^b1^−/−^ mice and the Cyp26b1^−/−^ mice in which the knockout was induced as adults also had significantly greater scores for otitis media, and half of the Cyp26b1^−/−^ mice in which knockout was induced as juveniles also showed otitis media (Figure 4 and Supplemental Figure 5). Increased keratin accumulation in the external ear canal was observed in all Cyp26a1^−/−^b1^−/−^ and Cyp26b1^−/−^ mice. Despite the greater survival of Cyp26b1^−/−^ mice in comparison to the Cyp26a1^−/−^b1^−/−^ mice, the histological findings in the skin and most of the severity scores appeared similar in-between the knockout models.

**Figure 4.**
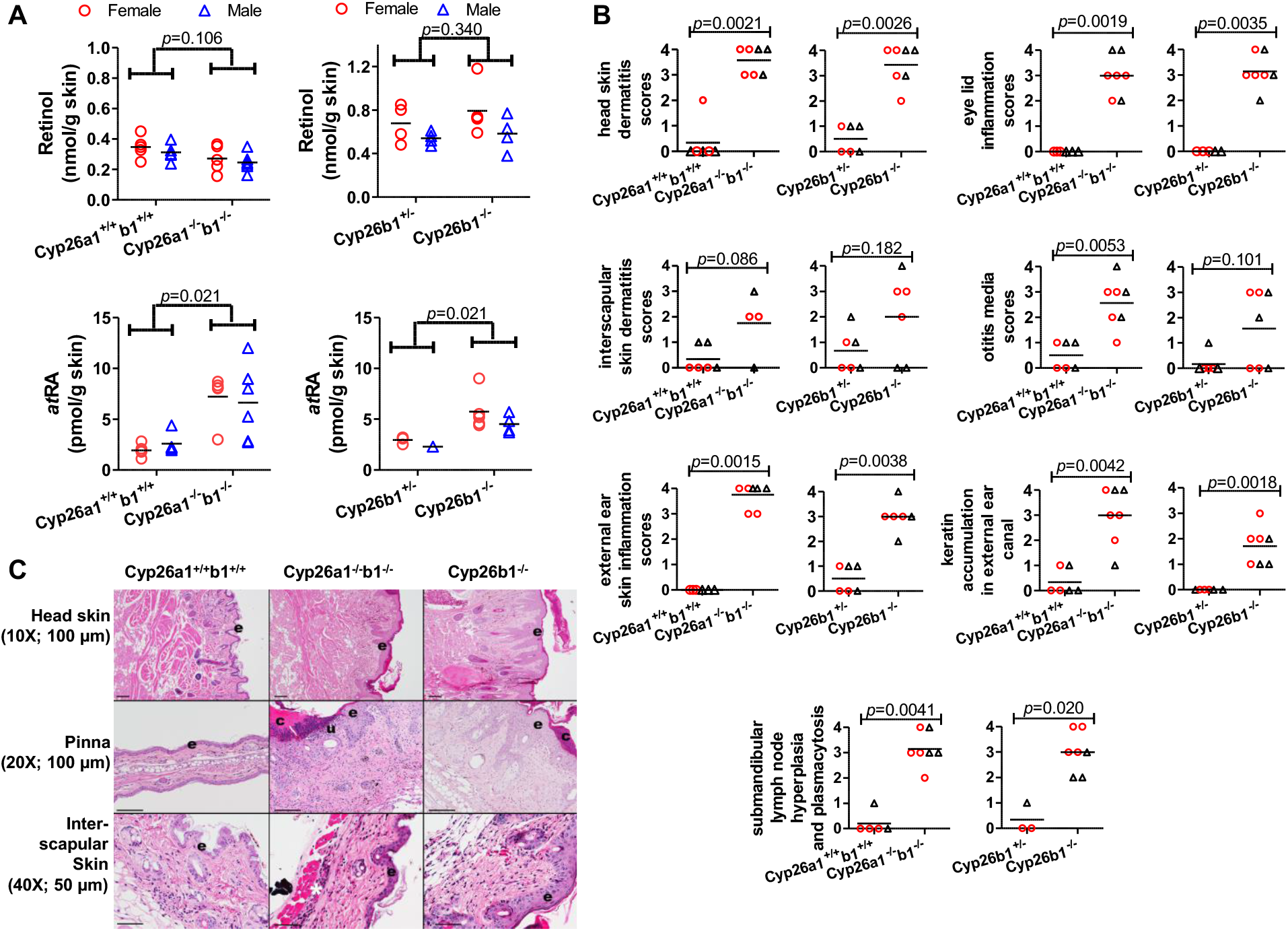
Skin retinol and *at*RA concentrations (A), histological scores (B) and H&E stained representative images (C) in mice in which knockouts were induced as juveniles. All the data in panel A are from mice in cohorts 2 and 3 and data in panels B and C are from mice in cohorts 1 and 2. In panels A and B, each data point represents an individual mouse. The horizontal lines indicate mean values. In panel B open red circles represent female mice and open black triangles show male mice. The differences between groups were tested by multivariate linear regression (A) and the Mann-Whitney test (B) and the obtained *p-*values are shown. In panel C, the original image magnification and the length that black scale bars represent are shown inside parentheses. “**e”**indicates epidermis; “**c”**indicates serocellular crust formation; “**u”**indicates epidermal erosion/ulceration; and asterisk indicates inflammation. The corresponding data for mice in which the knockout was induced as adults is shown in Supplemental Figure 5.

**Figure 5.**
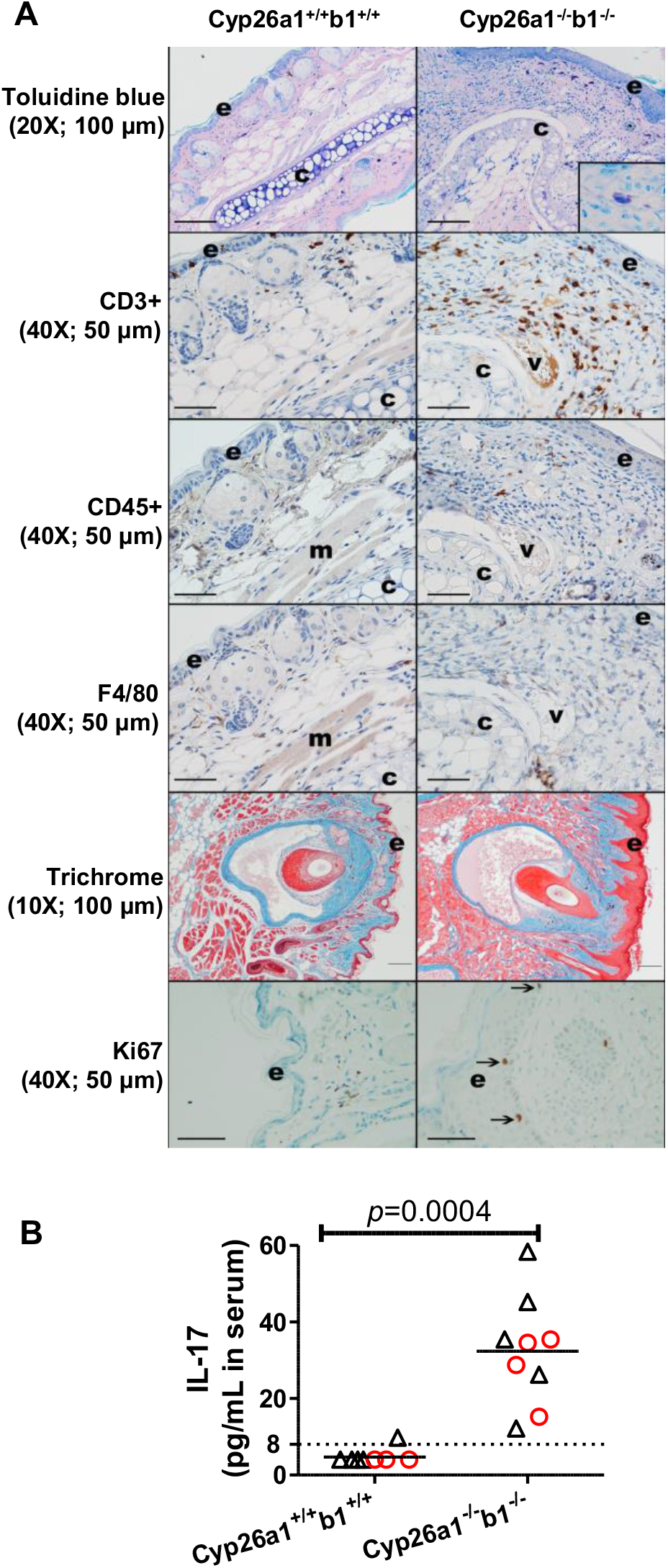
Representative skin images (A) and serum IL17 concentrations (B). All the data are from the juvenile mice in cohort 1. In panel A, the inset in the top right image shows a mast cell stained with toluidine blue. Arrows in the bottom panels indicate Ki67 positive cells in the stratum basale layer of the epidermis (e); “c” denotes cartilage; “v”, vessel and “m”, skeletal muscle. For CD45, CD3, F4/80 and Ki67 immunohistochemistry, immunoreactive cells are brown; blue = hematoxylin counterstain. Mild nonspecific staining is present in the vessel and skeletal muscle, and some pigment (melanin) is present in the skin. The original image magnification and the length that black scale bars represent are shown inside parentheses. In panel B, each data point represents an individual mouse. Serum IL17 LLOQ (8 pg/mL serum) is indicated by a dashed line. IL17 concentrations were assigned as 4 pg/mL (LLOQ/2) for samples that were below LLOQ. The horizontal lines indicate mean values. Red circles and black triangles represent female and male mice, respectively. The statistical significance was tested by Mann-Whitney test.

Retinoid concentrations were measured in the interscapular skin from the knockout mice compared to controls to explore the impact of Cyp26a1^−/−^b1^−/−^ and Cyp26b1^−/−^ on skin retinoid homeostasis. *at*RA concentrations were significantly increased in the skin of the Cyp26a1^−/−^b1^−/−^ mice, regardless on whether the knockout was induced in juveniles or adults (Figure 4 and Supplemental Figure 5). *at*RA concentrations were also significantly increased in the Cyp26b1^−/−^ mice in which the knock-out was induced as juveniles but not in the mice in which the knock-out was induced as adults (Figure 4 and Supplemental Figure 5). Skin retinol concentrations were decreased in the Cyp26a1^−/−^b1^−/−^ mice in which the knock-out was induced as adults but interscapular skin retinol concentrations were not significantly changed in either group of Cyp26b1^−/−^ mice or in the Cyp26a1^−/−^b1^−/−^ mice in which the knock-out was induced as juveniles.

Compared to controls and Cyp26b1^+/−^ mice, Cyp26a1^−/−^b1^−/−^ and Cyp26b1^−/−^ mice also had marked lymphadenomegaly of the mandibular lymph nodes with lymphoid hyperplasia and plasmacytosis. Mesenteric and popliteal lymph nodes were less consistently and severely affected (data not shown) suggesting that the lymphadenomegaly was at least partially due to inflammation of the skin in the head, otitis media and drainage from the head to the mandibular lymph nodes. Sporadic, generally mild lymphocytic salivary gland adenitis was observed in mice from all genotypes and considered a background lesion. In addition to the observed inflammation in the skin, circulating concentrations of IL-17 were also significantly increased (Figure 5) consistent with the role of this cytokine in skin inflammation. To better characterize the inflammatory cell infiltrate within the skin, immunohistochemistry staining for various cell types in the pinna was conducted in Cyp26a1^−/−^b1^−/−^ mice in comparison to controls (n = 1 female and 1 male mouse of each genotype). Based on H&E images and immunohistochemistry for CD3 (T lymphocytes), CD45 (B lymphocytes), F4/80 (macrophages), and Toluidine Blue (to highlight mast cells), the inflammatory cell infiltrate in the pinna of Cyp26a1^−/−^b1^−/−^ mice consisted predominantly of increased T-lymphocytes and neutrophils, with mast cells also subjectively increased compared to control mice (Figure 5). Trichrome images of the head highlighted the increased thickness of the epidermis (epithelial hyperplasia) and suggested mildly increased collagen in the dermis. Immunohistochemistry for Ki67 (a proliferation marker) showed increased positive cells in the stratum basale of the epidermis in Cyp26a1^−/−^b1^−/−^ mice in comparison to controls (Figure 5).

The mucosal surface of other squamous epithelium lined organs, including the esophagus and nonglandular stomach also appeared hyperplastic and/or hyperkeratotic in Cyp26a1^−/−^b1^−/−^ and Cyp26b1^−/−^ mice (Figure 6). The scores for non-glandular stomach hyperkeratosis and hyperplasia were significantly increased in the Cyp26a1^−/−^b1^−/−^ and Cyp26b1^−/−^ mice regardless of when the knockout was induced (adult or juvenile) (Figure 6 and Supplemental Figure 6). Stomach inflammation scores were also significantly increased in the Cyp26a1^−/−^b1^−/−^ and Cyp26b1^−/−^ mice in which knockout was induced as juveniles (Figure 6) but the increased inflammation scores in the stomach were not significant in the mice in which knockout was induced as adults (Supplemental Figure 6). At the limiting ridge, intracellular hyaline droplets (homogenous globular eosinophilic material in the cytoplasm of mucosal epithelial cells) and less commonly extracellular eosinophilic crystals, consistent with previous reports of material reacting to the Ym1/Ym2 protein, a member of the chitinase family (Ward *et al.*, 2001; Renne *et al.*, 2009; Nolte *et al.*, 2016), were observed to a greater degree in the glandular stomach of the juvenile-induction Cyp26a1^−/−^b1^−/−^ and Cyp26b1^−/−^ mice (Figure 6). These mice also tended to have more severe neutrophilic gastritis (Figure 6 and Supplemental Figure 6). Similar extracellular eosinophilic crystals were observed in the nasal cavity of Cyp26a1^−/−^b1^−/−^ mice (7/7 juvenile induction; 3/6 adult induction), less commonly in Cyp26b1^−/−^ mice (2/7 juvenile induction; no adult induction), and were not observed in control mice. YM1 immunohistochemistry of the stomach and nasal cavity in a single Cyp26a1^−/−^b1^−/−^ mouse showed that the intracellular eosinophilic globules were immunoreactive for YM1 (Supplemental Figure 7).

**Figure 6.**
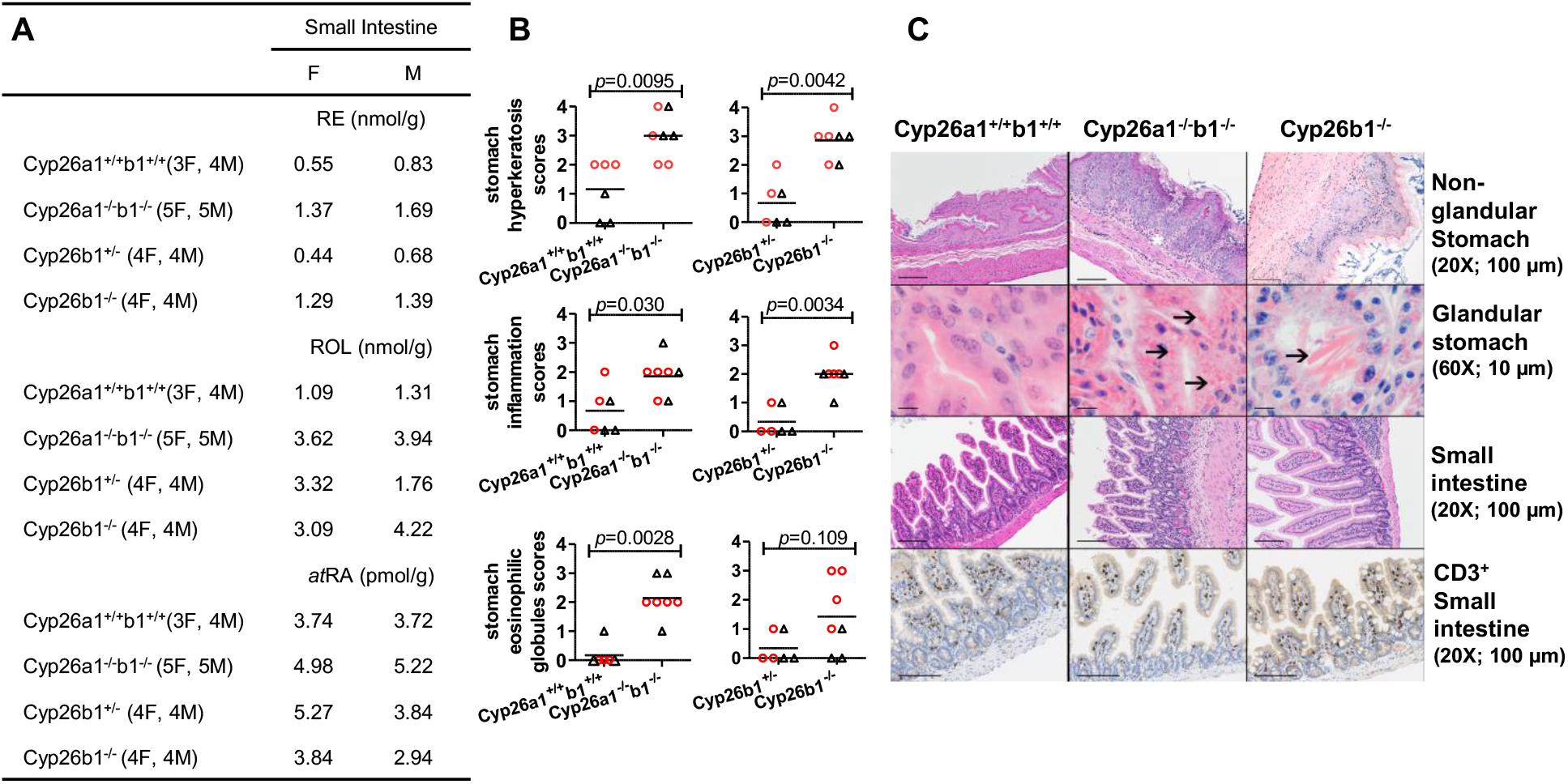
Concentrations of retinyl palmitate (RE), retinol (ROL) and *at*RA in the pooled mouse small intestine samples (A). The numbers of female, F, and male, M, mice from the indicated genotypes pooled in each group are shown inside parentheses and values for male and female are reported separately. The histological scores for the stomach hyperkeratosis, inflammation and hyaline droplets (eosinophilic globules) are shown in (B) and each data point represents an individual mouse with red circles and black triangles representing female and male mice, respectively. The horizontal lines indicate mean values and differences between groups were tested by the Mann Whitney test with *p*-values shown. H&E and anti-CD3 stained images of stomach and small intestine are shown in (C). All the data are from the juvenile mice in cohorts 1 and 2. In panel C, the original image magnification and the length black bars represent are shown inside parentheses. Intracellular hyaline droplets and extracellular eosinophilic crystals in the glandular stomach near the limiting ridge are indicated by arrows and the asterisk indicates inflammation in the nonglandular stomach.

On the large intestinal serosa, 3/6 juvenile induction Cyp26a1^−/−^b1^−/−^ mice had mild multifocal to coalescing chronic-active inflammation, thought to represent an extension of inflammation from the mesenteric fat, and 1 mouse had mild suppurative typhlocolitis with focal ulceration (data not shown). Routine histology did not show other consistent changes of the intestinal tract in Cyp26a1^−/−^b1^−/−^ and Cyp26b1^−/−^ mice, although some Cyp26a1^−/−^b1^−/−^ mice had villous atrophy and hypereosinophilic Paneth cells, most likely related to the poor nutritional status (Figure 6). To explore the phenotype further, Ki67 and CD3 immunohistochemistry was performed in the stomach and intestines of female juvenile-induction Cyp26a1^−/−^b1^−/−^ mice, but no obvious differences in percent positive Ki67 (stomach) or CD3 immunoreactivity (small intestine) were observed between Cyp26a1^−/−^b1^−/−^, Cyp26b1^−/−^ and control mice (Figure 6). Further investigation in a larger group of mice with more standardized sectioning (e.g. Swiss rolls or other preparation) might reveal differences not observed in the present study.

Retinoid concentrations in the intestines were analyzed in pooled samples from the mice, and the vitamin A metabolome in the small intestines appeared altered in both Cyp26a1^−/−^b1^−/−^ and Cyp26b1^−/−^ mice (Figure 6). Based on the analysis of 7-10 mice pooled per group, retinyl palmitate and retinol concentrations were two to three-fold higher in the Cyp26a1^−/−^b1^−/−^ and Cyp26b1^−/−^ mice than in the controls and *at*RA concentrations appeared elevated in particular in the Cyp26a1^−/−^b1^−/−^ mice.

### Postnatal global deletion of Cyp26a1 and Cyp26b1 or Cyp26b1 alone is associated with splenomegaly and increased myeloid:erythroid ratio in the bone marrow

During necropsy of the mice, moderate to severe splenomegaly was observed and the spleen weight in the Cyp26a1^−/−^b1^−/−^ and Cyp26b1^−/−^ mice was significantly increased (Figure 7 and Supplemental Figure 7). Compared to controls and Cyp26b1^+/−^ mice, Cyp26a1^−/−^b1^−/−^ and Cyp26b1^−/−^ mice had moderate expansion of the red pulp with increased myeloid and erythroid precursors and megakaryocytes (Figure 7 and Supplemental Figure 7). In sections of bone examined (decalcified sections of skull and femur), the myeloid:erythroid ratio appeared increased in the bone marrow of Cyp26a1^−/−^b1^−/−^ and Cyp26b1^−/−^ juvenile and adult induction mice (Supplemental Figure 7). As previous reports describe long-bone fractures and altered bone homeostasis in mice and rats suffering from *at*RA toxicity, H&E histology of the hind limb focused on the stifle with adjacent femur and tibia evaluated for adult induction Cyp26a1^−/−^b1^−/−^, Cyp26b1^−/−^ and control mice. In some mice, minimal tibia was available for review. Antemortem fractures were not noted histologically in the long bones of adult induction mice and no obvious consistent histologic differences of the femurs were noted between groups in adult induction mice, although the femurs appeared subjectively smaller in some Cyp26a1^−/−^b1^−/−^ mice compared to the other groups of mice.

**Figure 7.**
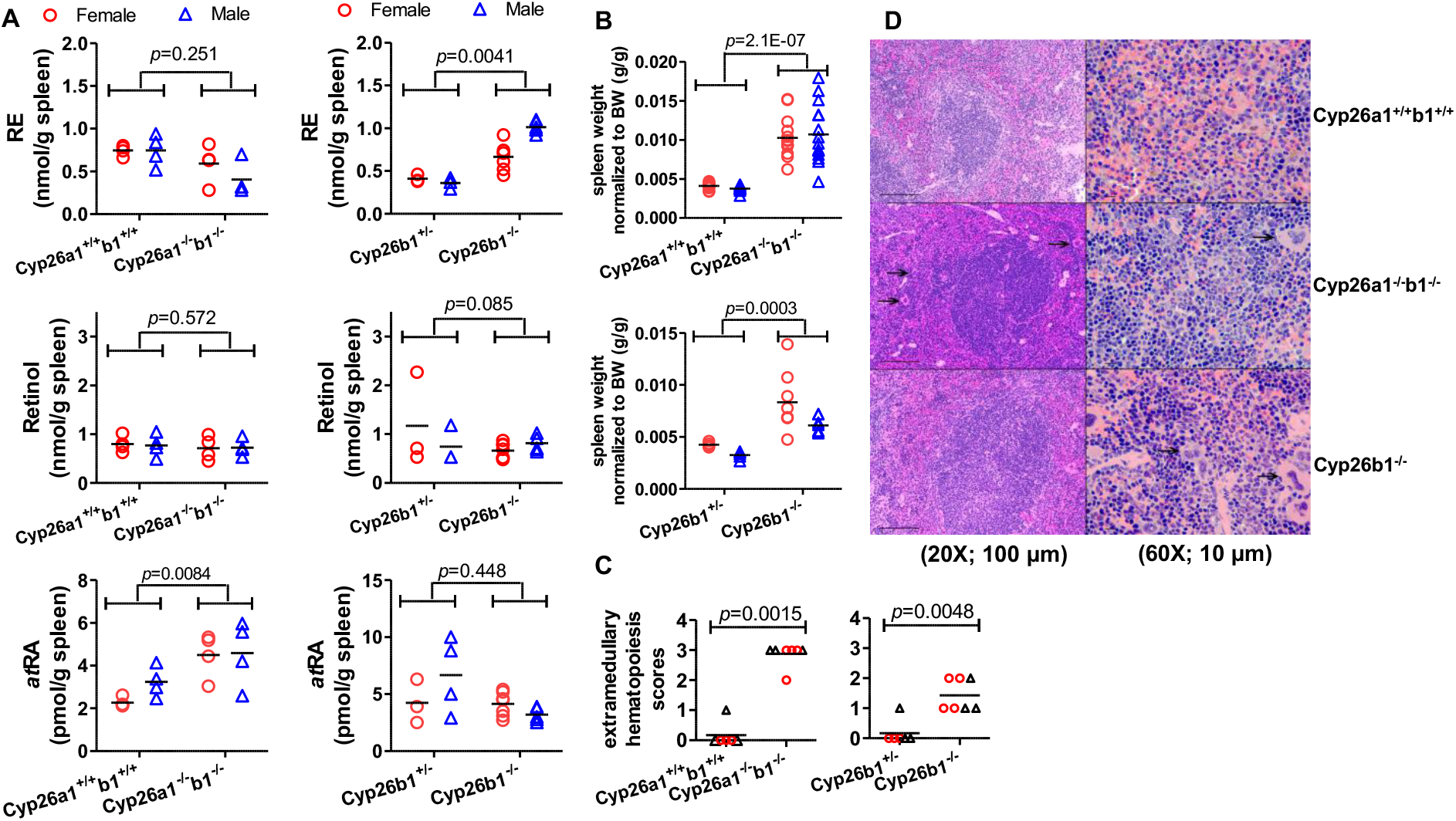
The concentrations of retinyl palmitate (RE), retinol and *at*RA in the spleen (A), spleen weight (B), histological scores of spleen hematopoietic cell proliferation (C), and H&E stained representative images (D). All the data are from the juvenile mice in cohorts 1 and 2. In A-C each data point represents an individual mouse and the horizontal lines indicate mean values. Differences between groups were tested by multivariate linear regression (A and B) or Mann Whitney test (C) and the obtained *p*-values are shown. The spleen weights were normalized to the terminal body weight (B). In panel C, red circles and black triangles represent female and male mice, respectively. In panel D, the original magnification and the length that black bars represent are shown inside parentheses. Arrows indicate megakaryocytes in the splenic red pulp. The corresponding data for mice in which the knockout was induced as adults is shown in Supplemental Figure 7.

Spleen concentrations of *at*RA were significantly increased in the Cyp26a1^−/−^b1^−/−^ mice when the knockout was induced as juveniles, while retinol and retinyl palmitate concentrations were largely similar across the groups (Figure 7 and Supplemental Figure 7). However, the total amount of the retinoids per spleen was significantly increased in the Cyp26a1^−/−^b1^−/−^ and Cyp26b1^−/−^ mice when compared to controls and Cyp26b1^+/−^ mice due to the increased size of the spleen (data not shown).

### Global deletion of Cyp26a1 and Cyp26b1 or Cyp26b1 alone is associated with inconsistent testicular degeneration

*at*RA is known to be critical for spermatogenesis, and regulation of *at*RA concentrations in the testis appears necessary for regulation of continuous spermatogenesis in the male. Male Cyp26a1^−/−^b1^−/−^ and Cyp26b1^−/−^ mice had variably severe testicular degeneration (Supplemental Figure 8). The testicular degeneration scores appeared similar in the adult induction Cyp26a1^−/−^b1^−/−^ and Cyp26b1^−/−^ mice while the testicular degeneration scores appeared elevated in the juvenile induction Cyp26a1^−/−^b1^−/−^ and to a lesser degree the Cyp26b1^−/−^ mice. No changes were observed in testicular retinoid concentrations with retinol, retinyl palmitate and *at*RA concentrations being similar in all groups of mice (Supplemental Figure 8).

### Clearance of exogenous atRA is significantly decreased in Cyp26a1^−/−^b1^−/−^ mice in comparison to control and these mice are hypersensitive to exogenous atRA

To test whether knockout of *Cyp26a1* and *Cyp26b1* alters the clearance of exogenous *at*RA, the PK of *at*RA was determined following a 1 mg/kg ip dose of *at*RA (Figure 8). The clearance of *at*RA was decreased by 80% (p<0.05) in the Cyp26a1^−/−^b1^−/−^ mice in comparison to controls, and the AUC of *at*RA increased by 5-fold (p<0.01) in the Cyp26a1^−/−^b1^−/−^ mice in comparison to controls (Figure 8). In addition to *at*RA, 13-*cis*RA was measured in all the samples, likely as an isomerization product from *at*RA. The AUC of 13-*cis*RA was also higher, by 3-fold, in the Cyp26a1^−/−^b1^−/−^ mice in comparison to controls.

**Figure 8.**
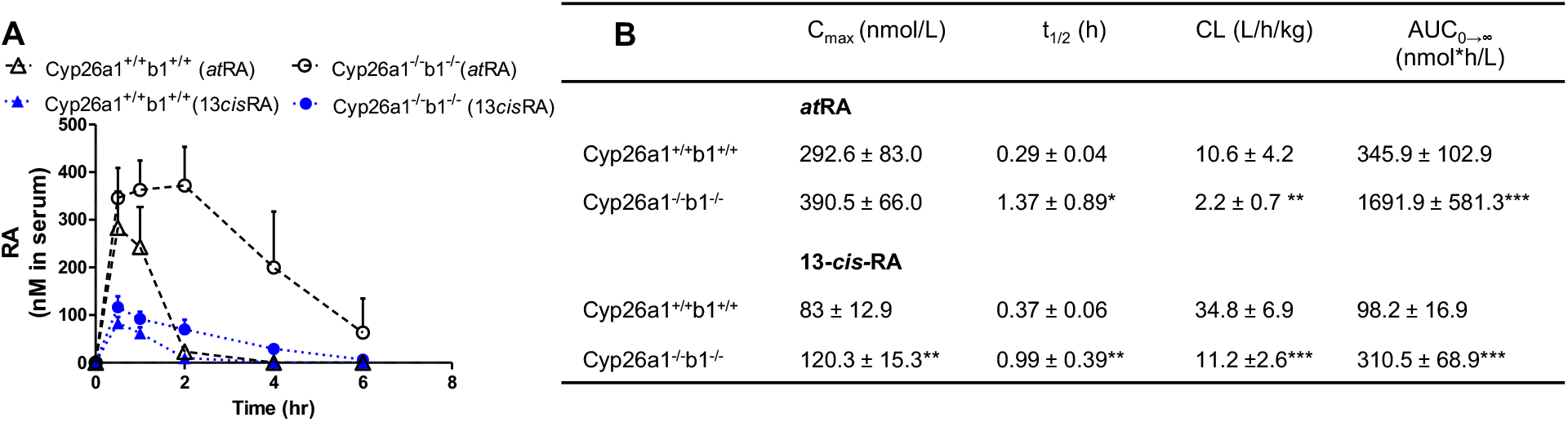
Pharmacokinetics (PK) of *at*RA (black symbols) and 13-*cis-*RA (blue symbols) following a 1 mg/kg ip injection to Cyp26a1+/+b1+/+ (control; triangles) and Cyp26a1−/−b1−/− (circles) mice. Panel A shows the plasma concentration-time data of *at*RA and 13-*cis*RA. Each data point indicates the mean value of retinoid concentrations measured (n=6) and error bars show SD. Panel B shows the PK parameters obtained. Data are shown as mean ± SD. Differences in PK parameters were tested by unpaired *t*-test. *, *p*<0.05; **, *p*<0.01; ***, *p*<0.001. All the data are from mice in cohort 4.

During the PK studies the behavior of the mice was also observed and the Cyp26a1^−/−^b1^−/−^ mice appeared to experience *at*RA toxicity. Following the dosing of *at*RA, the Cyp26a1^−/−^b1^−/−^ mice appeared weak and gathered together with decreased movement while control mice had normal behavior. At 4 hours after *at*RA dosing all of the Cyp26a1^−/−^b1^−/−^ mice had ruffled coats and still had diminished movement while the control mice did not appear to experience any toxicity. These observations followed the studies which initially dosed the mice with 10 mg/kg ip, the dose used in our previous studies without toxicities (Zhong *et al.*, 2019). The 10 mg/kg dose was not tolerated by the Cyp26a1^−/−^b1^−/−^ mice which experienced severe toxicity after this dose (90% mortality within 2 hours) while none of the control mice showed any signs of toxicity.

## Discussion

The absolute requirement for both Cyp26a1 and Cyp26b1 during fetal development has been thoroughly characterized (reviewed in (Pennimpede *et al.*, 2010)), and it is clear that these enzymes are critical in controlling *at*RA concentrations and signaling during embryonic development. While both Cyp26a1 and Cyp26b1 are required for normal development, the phenotypes of the global *Cyp26a1* and *Cyp26b1* knockout mice are distinct suggesting specific roles of the two enzymes. The phenotype of the *Cyp26a1* knockouts is more severe with hindbrain and neural tube defects, and the *Cyp26a1* knockout mice largely die during gestation (Abu-Abed *et al.*, 2001; Sakai *et al.*, 2001). *Cyp26b1* knockout mice die shortly after birth with respiratory distress and a variety of skeletal and limb defects (Yashiro *et al.*, 2004). Due to these severe phenotypes, very little is known about the specific roles of Cyp26a1 and Cyp26b1 in regulating retinoid signaling and concentrations in post-natal animals although it is generally accepted that at least Cyp26a1 functions as an enzyme that depletes bioactive retinoids rather than generates active retinoids (Niederreither *et al.*, 2002; Topletz *et al.*, 2015; Hernandez *et al.*, 2020). The expression patterns of Cyp26a1 and Cyp26b1 appear to be different in post-natal animals and during embryonic development with a clear shift in the expression patterns from fetal tissues to adult animals (Isoherranen and Zhong, 2019). For example, in post-natal animals Cyp26a1 appears to be the Cyp26 enzyme in the liver (Topletz *et al.*, 2012) while Cyp26b1 is the main *at*RA hydroxylase in bone marrow fibroblasts in vitro (Hernandez *et al.*, 2020). Recent studies of inducible global knockout of *Cyp26a1* in post-natal animals have shown that the roles of Cyp26 enzymes during fetal development cannot be extrapolated to post-natal life (Zhong *et al.*, 2019). The lack of an overt phenotype of the inducible *Cyp26a1* knockout mice together with kinetic modeling led us to hypothesize that in post-natal animals Cyp26b1 is, in fact, the main enzyme clearing endogenous *at*RA. Collectively the data in this study support the hypothesis that Cyp26b1 is the main post-natal *at*RA clearing enzyme. This study also emphasizes the importance of Cyp26a1 as an inducible backup *at*RA hydroxylase in post-natal animals that appears to contribute to the regulation of overall retinoid exposures and attenuates potential retinoid toxicities.

Postnatal global knockout of *Cyp26b1* alone or together with *Cyp26a1* resulted in severe dermatitis with epidermal hyperplasia and hyperkeratosis, blepharitis, splenomegaly, lymphadenomegaly and inflammation, giving the two knockout models a similar appearance. However, the Cyp26b1^−/−^ mice reproducibly showed higher *at*RA concentrations only in the skin. This suggests that Cyp26b1 is particularly relevant for regulating *at*RA concentrations and signaling in this organ. The observed dermatitis, blepharitis, and skin inflammation are likely due to the loss of Cyp26b1 mediated *at*RA clearance and subsequent localized increase in skin *at*RA concentrations. High expression levels of Cyp26b1 have previously been reported within skin fibroblasts, and *at*RA has been shown to trigger an inflammatory response via P2X7 activation in cultured skin fibroblasts (Kurashima *et al.*, 2014). When mice were treated with *at*RA and liarozole, an inhibitor of the Cyp26 enzymes, severe skin inflammation was observed in the ears (Kurashima *et al.*, 2014). In fact, the previous observations using liarozole were very similar to the current findings in Cyp26b1^−/−^ and Cyp26a1^−/−^b1^−/−^ mice. Consistent with the previous findings that *at*RA induces mast cell activation via P2X7 receptor activation, mast cells appeared to be increased in the skin of the Cyp26a1^−/−^b1^−/−^ mice in addition to other inflammatory cells such as neutrophils and T-lymphocytes. Overall, the results presented here suggest that Cyp26b1 is the main *at*RA hydroxylase in the skin, and that selective Cyp26b1 inhibitors may be beneficial over pan-Cyp26 inhibitors such as talarozole or liarozole for treatment of various skin conditions that are retinoid responsive. Splenic and bone marrow changes with increased hematopoietic precursors were considered likely to be at least partially related to the severe dermatitis (Hampton *et al.*, 2015).

The role of Cyp26b1 in comparison to Cyp26a1 as *at*RA hydroxylase in other organs except the skin is less clear based on the data collected, and it appears that both Cyp26a1 and Cyp26b1 contribute to regulation of *at*RA concentrations in post-natal animals either via built-in redundancies or by compensating for each other. While *at*RA concentrations were unaltered in the Cyp26a1^−/−^ mice described previously (Zhong *et al.*, 2019) and in the Cyp26b1^−/−^ mice described here (except the skin), *at*RA concentrations in serum and in various tissues were overall increased in the Cyp26a1^−/−^b1^−/−^ mice. The fact that *at*RA concentrations were increased in Cyp26a1^−/−^b1^−/−^ mice overall supports the critical role the two Cyp26 enzymes play together in clearing *at*RA. The phenotype of the Cyp26a1^−/−^b1^−/−^ mice was generally more severe than either Cyp26a1^−/−^ (Zhong *et al.*, 2019) or Cyp26b1^−/−^ mice. For example, Cyp26b1^−/−^ mice gained weight similarly to control or Cyp26b1^+/−^ mice while Cyp26a1^−/−^b1^−/−^ mice suffered from weight loss or did not gain as much weight as controls and also had more pronounced fat inflammation and adipose tissue atrophy observed histologically. In addition, while all Cyp26b1^−/−^ mice survived till the end of the observation period, almost half of the juvenile male and female and half of the adult female induction Cyp26a1^−/−^b1^−/−^ mice had to be sacrificed early. These findings highlight the critical role that Cyp26a1 plays in tissue retinoid homeostasis despite the lack of phenotype of post-natal *Cyp26a1* single knockout mice (Zhong *et al.*, 2019).

The phenotype observed in the Cyp26a1^−/−^b1^−/−^ mice is very similar to the observed phenotype in mice and rats after chronic dosing with *at*RA. While acute *at*RA toxicity results in hair loss, dermal and mucosal alterations and loss of body weight (Kretzschmar and Leuschner, 1975; Biesalski, 1989), chronic *at*RA treatment has been shown to cause lymphoid hyperplasia, extramedullary hematopoiesis and hyperkeratosis (Kurtz *et al.*, 1984), findings that are very similar to the phenotype observed in this study. The observations that exogenous *at*RA clearance was reduced in the Cyp26a1^−/−^b1^−/−^ mice by 80% and the Cyp26a1^−/−^b1^−/−^ mice were hypersensitive to *at*RA toxicity, further confirm the critical role of Cyp26a1 and Cyp26b1 in clearing *at*RA in post-natal animals. Previous reports have described long bone fractures and altered bone homeostasis in mice and rats treated with *at*RA. We did not observe fractures or obvious histologic abnormalities other than a smaller appearance to the femur in adult induction dual knockout mice, although we did not dissect out the bones for gross examination or perform radiography or other imaging. Knockout of *Cyp26b1* or mutations of *Cyp26b1* have been previously shown to result in variety of bone abnormalities. Ablation of *Cyp26b1* in chondrocytes induced a gradual decrease in bone length beginning two weeks after birth and premature closing of the growth plate (Minegishi *et al.*, 2014). Similarly, disruption of *Cyp26b1* has been shown to alter ossification and osteoblast activity (Laue *et al.*, 2008; Spoorendonk *et al.*, 2008) and mutations in *Cyp26b1* generally affect bone growth (Isoherranen and Zhong, 2019). It is possible that the lack of a robust bone phenotype in this study is due to the timing of the induction of the knockout, and that *Cyp26b1* must be knocked out either during fetal development or during earlier post-natal development to result in dramatic skeletal abnormalities. Further workup in both juvenile and adult induction mice is warranted to further characterize a possible, more subtle phenotype, particularly as it relates to osteoblast and osteoclast activity.

Similar to the bone, it is likely that Cyp26b1 plays a critical role in testis development during fetal development and early post-natal life, and the timing of when the knockout is induced is critical for observed testis phenotype. In a previous study, simultaneous deletion of *Cyp26b1* in Sertoli and germ cells resulted in male subfertility and loss of advanced germ cells from the seminiferous epithelium (Hogarth *et al.*, 2015). In this study the testicular degeneration, albeit sporadic, was more severe in the mice in which the knockout was induced as juveniles and was most clearly observed in the Cyp26a1^−/−^b1^−/−^ mice when knockout was induced as juveniles. Yet, the phenotype observed in these mice was less severe than that observed in the mice with deletion of *Cyp26b1* in Sertoli and germ cells (Hogarth *et al.*, 2015) and required knockout of both *Cyp26a1* and *Cyp26b1*. This suggests that Cyp26b1 plays an important role in regulating *at*RA concentrations during the initiation of spermatogenesis, but once spermatogenesis is established the testis becomes less sensitive to the loss of the Cyp26 enzymes.

One of the challenges in defining the specific roles of *at*RA versus its main circulating precursor, retinol and the binding protein RBP4 is the lack of tools that would allow modulating solely *at*RA concentration or retinol-RBP4 without altering the other. Disrupting either *at*RA concentrations or retinol-RBP4 levels typically results in alterations in the other due to the feedback loops that control retinoid homeostasis. This challenge was clearly shown in this study as well. The Cyp26a1^−/−^b1^−/−^ mice not only had increased *at*RA concentrations in affected tissues but they also showed a decrease in retinol and RBP4 concentrations in serum and increased retinol and retinyl ester concentrations in the small intestine. It is possible that some of the lesions observed in Cyp26a1^−/−^b1^−/−^ mice are in fact due to the decrease in retinol and RBP4 in circulation rather than increased *at*RA, and this requires further study. In this knockout model, there is a disconnect between low levels of retinol and RBP4 in the serum and the high concentrations of *at*RA. While not surprising given the tightly regulated feedback loops that control retinoid homeostasis, these observations come to highlight the limitations of using serum retinol and RBP4 levels as surrogate markers of tissue *at*RA and vitamin A concentrations.

In conclusion, this study was designed to test the hypothesis that Cyp26 enzymes and Cyp26b1 in particular are critical for *at*RA clearance in post-natal animals and regulate endogenous *at*RA concentrations and signaling. This study shows that Cyp26 enzymes are necessary for post-natal life and lack of these enzymes leads to retinoid toxicity characterized by dermatitis, hyperkeratosis and hyperplasia of squamous epithelia and inflammation. Cyp26b1 alone was found to be critical for maintaining retinoid homeostasis in the skin while both Cyp26 enzymes appeared necessary for overall retinoid homeostasis. This study also suggests that while the liver plays a critical role in storing retinyl esters and in synthesizing RBP4 this organ is relatively resistant to short term *at*RA toxicity. The findings presented here lay the foundation for future studies of the role of Cyp26 enzymes in gut homeostasis, immune regulation, inflammation, hematopoiesis and bone homeostasis.

## Acknowledgments and Funding

This study was supported by an NIH grant R01 GM111772 (to JS, CH and NI) and T32 DK007247 (to LC)

The authors wish to thank Brian Johnson and Histology and Imaging Core for technical expertise with histology and immunohistochemistry.

