## Supplemental materials, figures and tables for "Knockout of Cyp26a1 and Cyp26b1 during post-natal life causes reduced lifespan, dermatitis, splenomegaly and systemic inflammation in mice"

**Histology semi-quantitative scoring criteria:**

Skin (decalcified cross sections of skull); pinna

1: Minimal to mild focal or bilateral (muzzle only) inflammation +/- hyperkeratosis; 2: Mild to moderate multifocal to coalescing acanthosis and hyperkeratosis +/- mild inflammation; 3: Moderate multifocal to coalescing acanthosis, hyperkeratosis, and inflammation with epidermal erosion; 4: Severe acanthosis, hyperkeratosis, and inflammation with ulceration or multifocal erosion and serocellular crusting

Blepharitis:

1: Minimal bilateral or mild unilateral; 2: Mild bilateral (acanthosis/hyperkeratosis with mild inflammation); 3: Moderate bilateral; 4: Severe bilateral with ulceration and extensive inflammation

Otitis media:

1: Mild unilateral proteinaceous fluid with minimal-mild cellular component; 2: Moderate-severe inflammation and proteinaceous fluid, unilateral; 3: Mild bilateral; 4: Moderate-severe bilateral

External ear canal keratin accumulation:

1: Mild unilateral or minimal bilateral; 2: Moderate-severe unilateral; 3: Mild-moderate bilateral; 4: Severe bilateral (occluding ear canal)

Testicular degeneration:

1: Minimal - occasional tubular vacuolation or multinucleated germ cells; 2: Mild - tubular vacuolation and germ cell degeneration/depletion affecting 5-25%; of testis 3: Moderate – affects 25-65% of testis; 4: Severe, >66% of testis with loss of normal architecture

Hyperkeratosis and inflammation, nonglandular stomach:

1: Minimal; 2: Mild, involving <50% of nonglandular stomach with keratin retention and scattered inflammation; 3: >50-75% of nonglandular stomach to any degree or focally extensive moderate keratin retention and mild-moderate inflammation; 4: Severe, involving >75% nonglandular stomach

Eosinophilic globules, glandular stomach:

1: Minimal, scattered in region of limiting ridge only; 2: Mild, generally adjacent to limiting ridge; 3: Moderate adjacent to limiting ridge, extensive along 30-50% of glandular stomach, larger; 4: Severe, extensive along >50% of glandular stomach

Lymphocyte/plasma cell hyperplasia (lymph node):

1: Minimal lymphocyte or plasma cell hyperplasia; 2: mildly enlarged, increased plasma cells; 3: moderately enlarged, >50% plasma cells; 4: markedly enlarged, >66% plasma cells +/- neutrophils, histiocytes

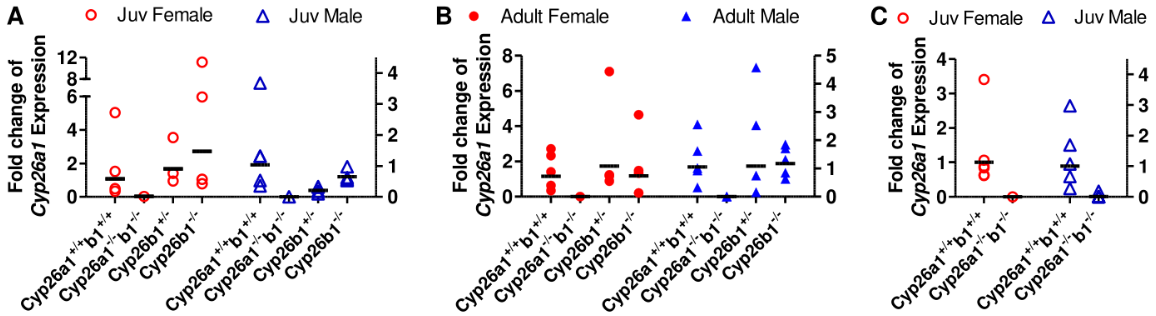


Supplemental Figure 1. The relative Cyp26a1 mRNA expression in mouse liver. Each data point represents an individual mouse. The horizontal lines indicate geometric mean values for the group. The fold change is the average value calculated from two rt-PCR analyses performed on separate days. Female and male data were analyzed separately and represented on the left and right Y axis, respectively. Data in panels A and B are from mice in cohorts 1 and 2 while data in panel C is from cohort 3.

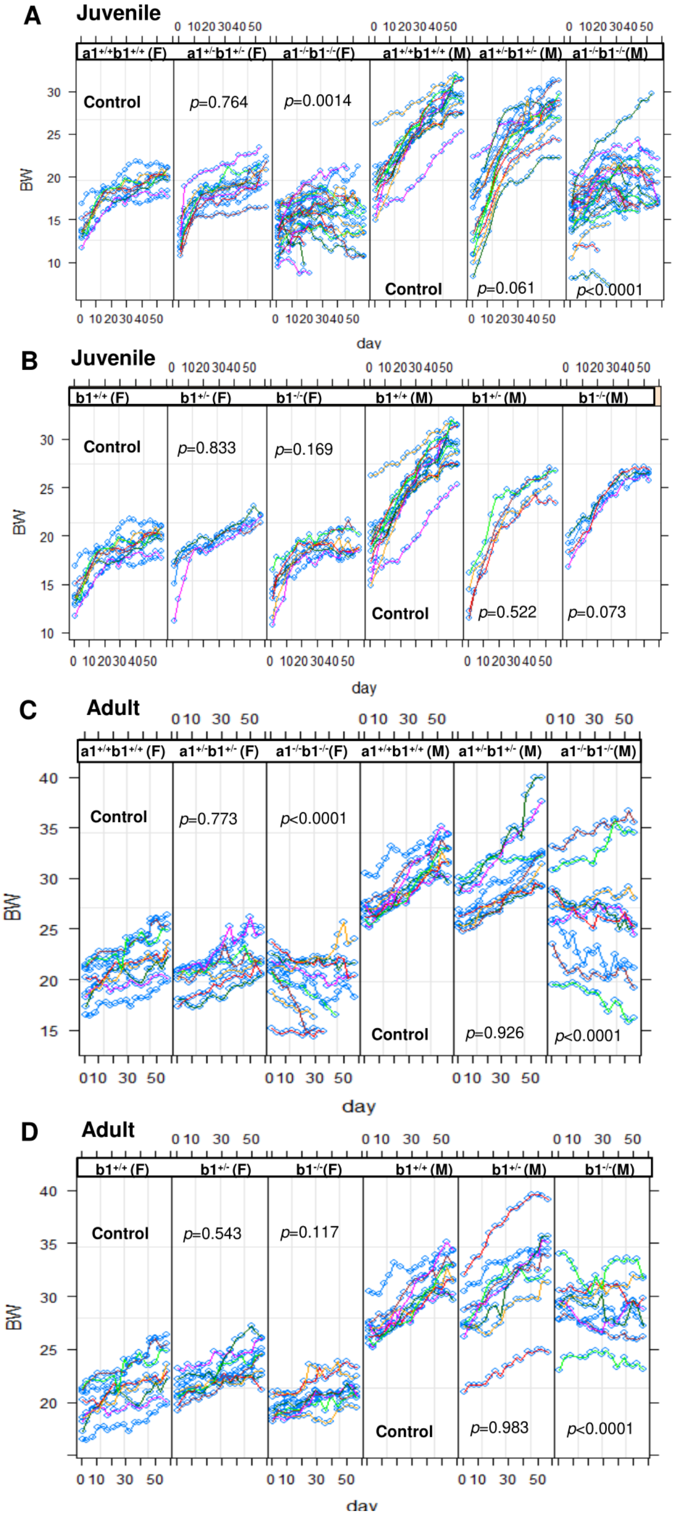


Supplemental Figure 2. Body weight analyses by the multivariate linear mixed effect model. The X axis shows days after tamoxifen injection. All the data are from mice in cohorts 1 and 2. The *p* values shown in the figure indicate the statistical difference of the slope in comparison to the control mice. F, female; M, male. $a1^{+/+}b1^{+/+}$, $Cyp26a1^{+/+}b1^{+/+}$; $a1^{+/-}b1^{+/-}$, $Cyp26a1^{+/-}b1^{+/-}$; $a1^{-/-}b1^{-/-}$, $Cyp26a1^{-/-}b1^{-/-}$; $b1^{+/+}$, $Cyp26b1^{+/+}$; $b1^{+/-}$, $Cyp26b1^{+/-}$; $b1^{-/-}$, $Cyp26b1^{-/-}$.

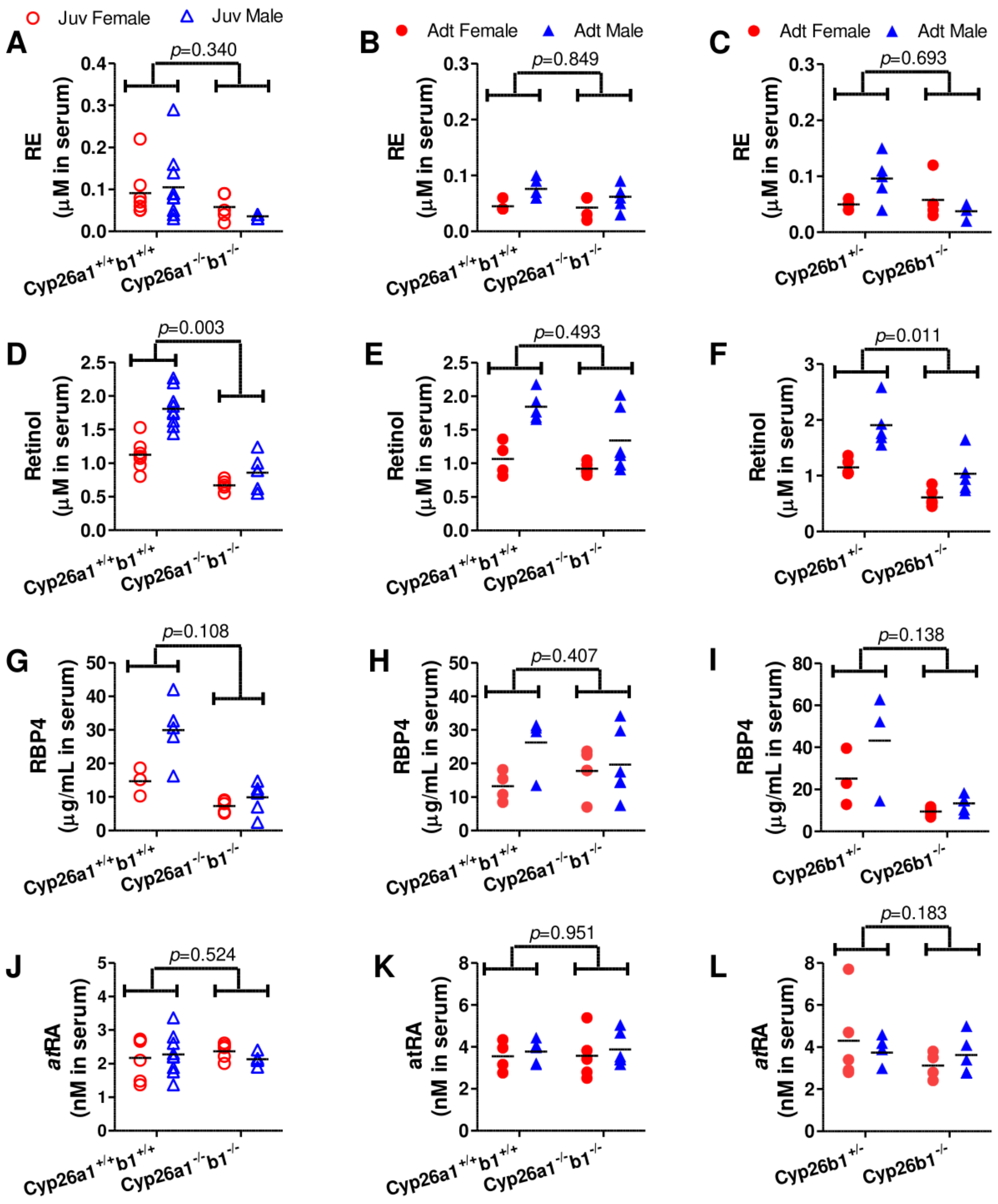


Supplemental Figure 3. Serum retinoid and RBP4 concentrations in Cyp26a1$^{+/+}$b1$^{+/+}$, Cyp26a1$^{-/-}$b1$^{-/-}$, Cyp26b1$^{+/-}$ and Cyp26b1$^{-/-}$ mice. All the data are from mice in cohorts 1 and 2. Each data point represents an individual mouse. The horizontal lines indicate mean values. The *p* values were obtained from statistical analyses done by multivariate linear regression. Juv, juvenile; Adt, adult.

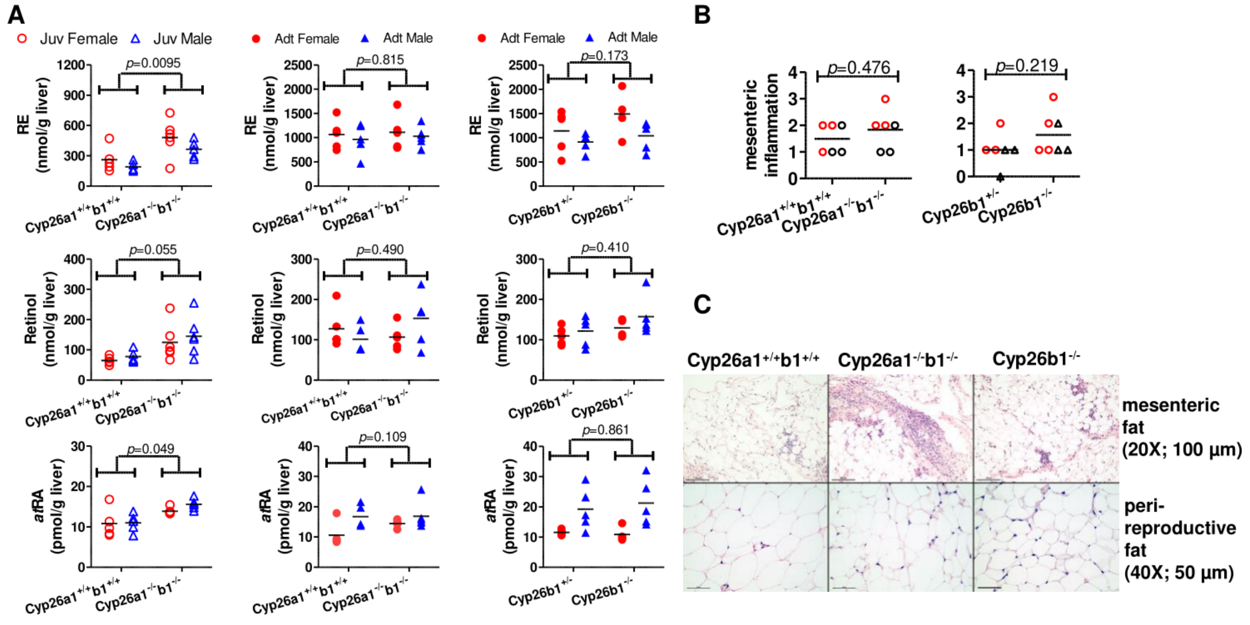


Supplemental Figure 4. Liver retinoid concentrations (A), histological scores of mesenteric inflammation in adipose (B) and H&E stained representative images of adipose (C). All the data are from mice in cohorts 1 and 2. Each data point represents an individual mouse. The horizontal lines indicate mean values. The *p* values were obtained from statistical analyses analyzed by multivariate linear regression (A) and Mann Whitney test (B). In panel B, red circles and black triangles represent female and male mice, respectively. In panel C, the original image magnification and the length of the black scale bars are listed inside parentheses. Juv, juvenile; Adt, adult.

Supplemental Table 1. Summary of p values for testing for sex differences in the Cyp26a1$^{+/+}$b1$^{+/+}$ and Cyp26a1$^{-/-}$b1$^{-/-}$ mice.

| | | RE | ROL | RBP4 | *at*RA |
|---|---|---|---|---|---|
| serum | Juv-3 | 0.471 | **0.035** | 0.083 | 0.100 |
| | Juv | 0.643 | **2.15E$^{-06}$** | **0.0006** | 0.699 |
| | Adt | **0.023** | **0.002** | **0.036** | 0.936 |
| liver | Juv-3 | 0.688 | 0.437 | N/A | 0.410 |
| | Juv | 0.380 | 0.667 | N/A | 0.895 |
| | Adt | 0.601 | 0.389 | N/A | **0.016** |
| skin | Juv-3 | N/A | 0.425 | N/A | 0.653 |
| | Adt | N/A | 0.002 | N/A | 0.901 |
| spleen | Juv-3 | 1.000 | 0.854 | N/A | 0.196 |
| | Adt | 0.099 | 0.483 | N/A | 0.112 |

Note: Juv, juvenile; Adt, adult. Juv-3 means the juvenile mice from cohort 3 while Juv and Adt indicate juvenile and adult mice from groups from cohorts 1 and 2.

Supplemental Table 2. Summary of p values for testing for sex differences in the Cyp26b1$^{+/-}$ and Cyp26b1$^{-/-}$ mice.

| | | RE | ROL | RBP4 | *at*RA |
|---|---|---|---|---|---|
| serum | Juv | 0.421 | **0.0001** | **0.0004** | 0.153 |
| | Adt | **0.036** | **0.0009** | 0.051 | 0.376 |
| liver | Juv | 0.628 | 0.861 | N/A | 0.980 |
| | Adt | 0.415 | 0.590 | N/A | **0.042** |
| skin | Juv | N/A | 0.280 | N/A | 0.698 |
| | Adt | N/A | 0.089 | N/A | 0.105 |
| spleen | Juv | 0.585 | 0.213 | N/A | 0.109 |
| | Adt | 0.139 | 0.018 | N/A | 0.697 |

Note: Juv, juvenile; Adt, adult. Data were obtained from mice in cohorts 1 and 2.

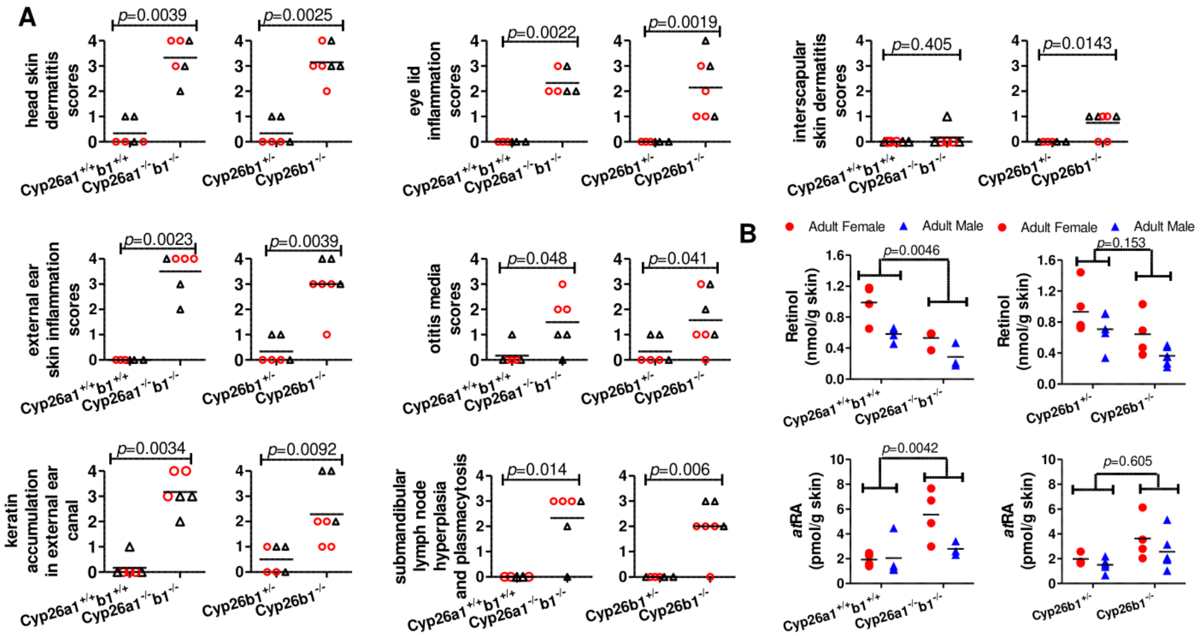


Supplemental Figure 5. Skin histological scores (A) and retinoid concentrations (B). All the data are from the adult mice in cohorts 1 and 2. Each data point represents an individual mouse. The horizontal lines indicate mean values. In panel A, red circles and black triangles represent female and male mice, respectively. The statistical significance was analyzed by Mann-Whitney test (A) and multivariate linear regression (B).

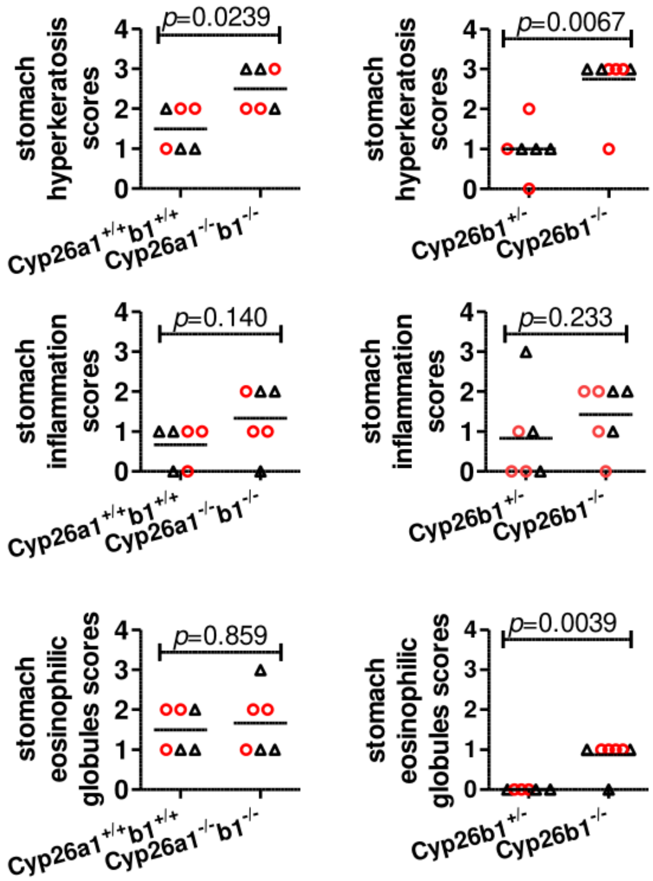


Supplemental Figure 6. Stomach histological scores. All the data are from the adult mice in cohorts 1 and 2. Each data point represents an individual mouse. Red circles and black triangles represent female and male mice, respectively. The horizontal lines indicate mean values. The statistical significance was analyzed by Mann Whitney test.

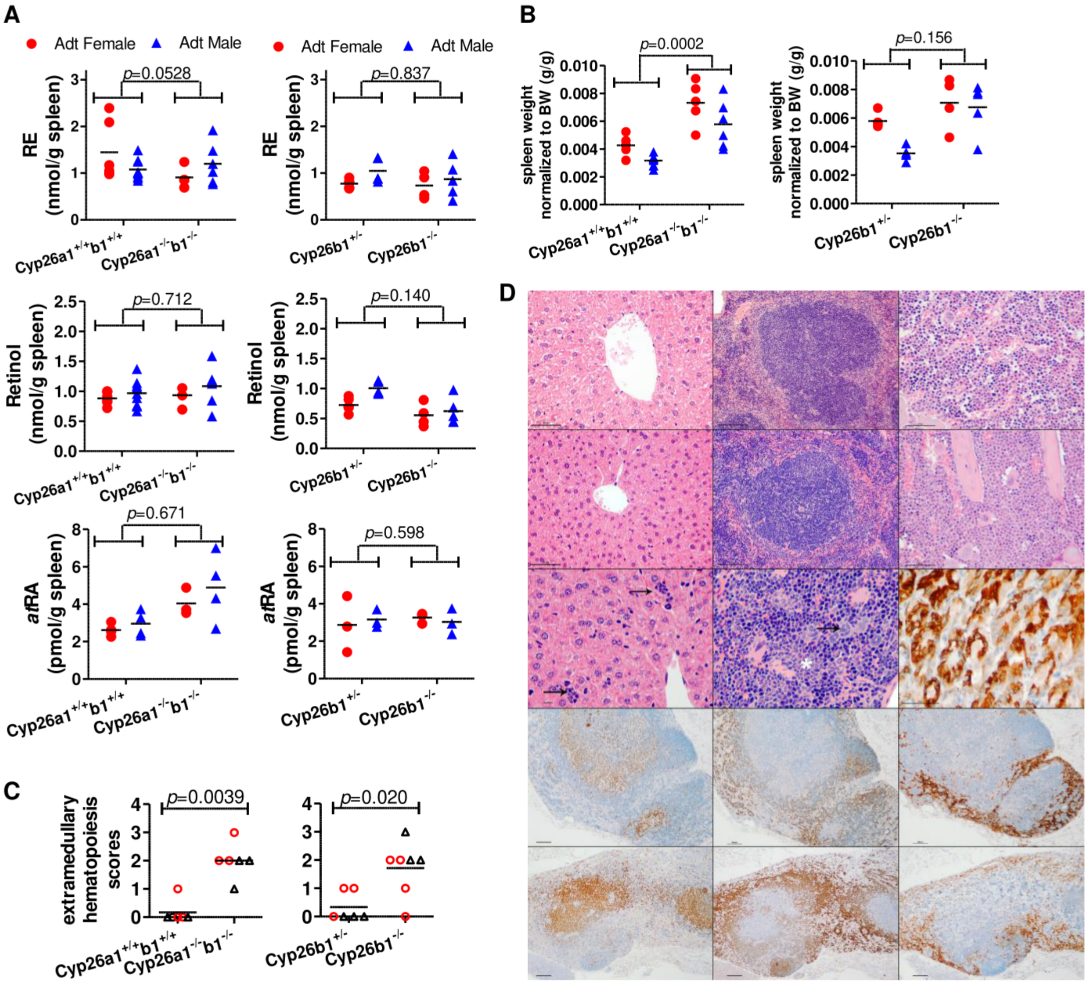


Supplemental Figure 7. Spleen retinoid concentrations (A), spleen weight (B), histological scores of spleen hematopoietic cell proliferation (C) and H&E and IHC stained representative images of liver, spleen, bone marrow and lymph node (D). All the data are from mice in cohorts 1 and 2. In panels A to C, each data point represents an individual mouse. The horizontal lines indicate mean values. The *p* values were obtained from statistical analyses done by multivariate linear regression (A and B) and Mann Whitney test (C). The spleen weight was normalized to the terminal body weight (B). Red circles and black triangles represent female and male mice, respectively (C). In panel D, the top row includes H&E images from Cyp26a1$^{+/+}$b1$^{+/+}$ mice; from left to right are liver from an adult male mouse (40X, bar-50 μm), spleen from a juvenile male mouse (20X, bar-100 μm) and bone marrow from a juvenile female mice (40X, bar-50 μm). The 2$^{nd}$ row includes H&E images; from left to right are liver from a Cyp26a1$^{-/-}$b1$^{-/-}$ juvenile male (40X, bar-50 μm), spleen from a Cyp26a1$^{-/-}$b1$^{-/-}$ juvenile male (20X, bar-100 μm) and bone marrow from a Cyp26b1$^{-/-}$ juvenile female (40X, bar-50 μm); the bone marrow H&E image from a Cyp26b1$^{-/-}$ juvenile female shows an increased ratio of myeloid to erythroid cells in comparison to the control group (Cyp26a1$^{+/+}$b1$^{+/+}$). The 3$^{rd}$ row includes H&E and IHC images; from left to right are liver from a Cyp26a1$^{-/-}$b1$^{-/-}$ adult female (60X, bar-10 μm), spleen from a Cyp26a1$^{-/-}$b1$^{-/-}$ adult female (60X, bar-10 μm) and anti-YM1 IHC image of stomach from a Cyp26a1$^{-/-}$b1$^{-/-}$ juvenile male (40X, bar-50 μm); arrows in the liver H&E image indicate extramedullary proliferation of hematopoietic precursor cells; the arrow in the spleen H&E image indicates megakaryocytes and the asterisk indicates myeloid precursors. The 4$^{th}$ and 5$^{th}$ rows include representative IHC stained images of mesenteric lymph node from juvenile mice in the Cyp26a1$^{+/+}$b1$^{+/+}$ (4$^{th}$ row) and Cyp26a1$^{-/-}$b1$^{-/-}$ (5$^{th}$ row) group; from left to right are CD3, CD45 and F4/80 images; the original magnification is10X and bars represent 100 μm.

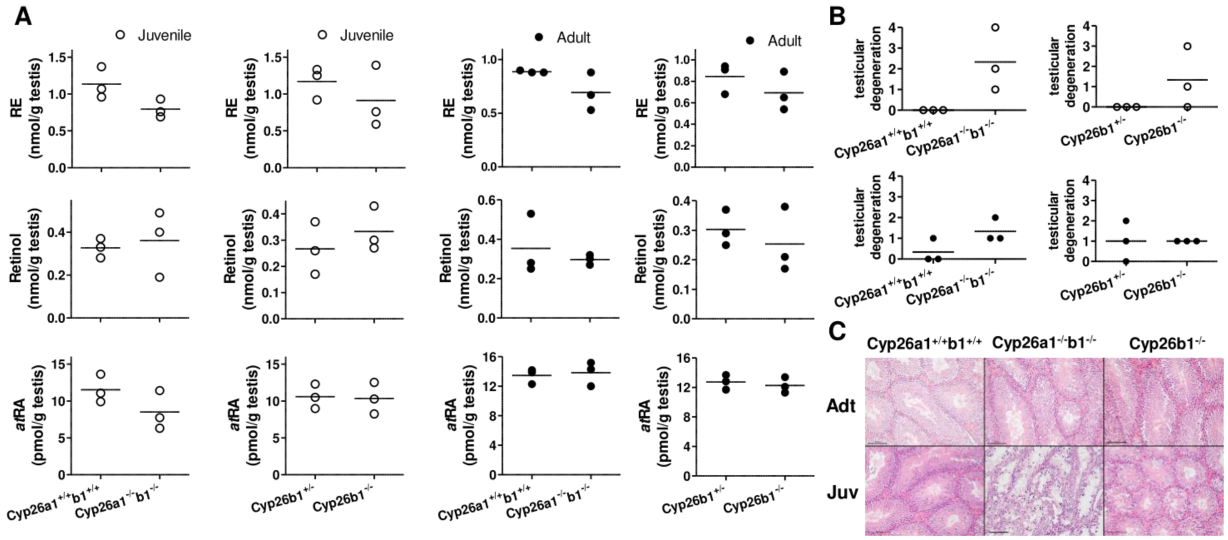


Supplemental Figure 8. Testicular retinoid concentrations (A), testicular degeneration scores (B) and H&E stained representative images of testis (C). The statistical analyses were done using unpaired *t*-test (A) and Mann Whitney test (B). All the *p* values were greater than 0.05. In panel C the magnification is 20X and black bars in the images represent 100 µm. All the data are from mice in cohorts 1 and 2.
